# Direct detection of 8-oxo-dG using nanopore sequencing

**DOI:** 10.1101/2024.05.17.594638

**Authors:** Marc Pagès-Gallego, Daan M. K. van Soest, Nicolle J. M. Besselink, Roy Straver, Janneke P. Keijer, Carlo Vermeulen, Alessio Marcozzi, Markus J. van Roosmalen, Ruben van Boxtel, Boudewijn M. T. Burgering, Tobias B. Dansen, Jeroen de Ridder

## Abstract

Genomic DNA is constantly subjected to oxidative damage, which is thought to be one of the major drivers of cancer and age-dependent decline. The most prominent consequence is the modification of guanine into 8-hydroxyguanine (8-oxo-dG), which has important mutagenic potential and plays a role in methylation-mediated gene regulation. Methods to simultaneously detect and quantify 8-oxo-dG within its genomic context have been lacking; mainly because these methods rely on indirect detection or are based on hydrolysis of the DNA. Nanopore sequencing has been deployed for the direct detection of base-modifications like cytosine methylation during sequencing. However, currently there is no model to detect 8-oxo-dG by nanopore sequencing due to the lack of training data. Here, we developed a strategy based on synthetic oligos to create long DNA molecules with context variability for effective deep learning and nanopore sequencing. Moreover, we showcase a training approach suitable to deal with the extreme scarceness of 8-oxo-dG compared to canonical G to enable specific 8-oxo-dG detection. Applied to an inducible tissue culture system for oxidative DNA damage, our approach reveals variable 8-oxo-dG distribution across the genome, a dissimilar context pattern to C>A mutations, and concurrent 5-mC depletion within a 2-kilobase window surrounding 8-oxo-dG sites. These findings not only underscore the potential of nanopore sequencing in epigenetic research, but also shed light on 8-oxo-dG’s role in genomic regulation. By simultaneously measuring 5-mC and 8-oxo-dG at single molecule resolution, our study provides insights into the functional interplay between these DNA modifications. Moreover, our approach using synthetic oligos to generate a ground truth from machine learning modification calling could be applied to any other DNA modification. Overall, our work contributes to advancing the field of epigenetics and highlights nanopore sequencing as a powerful tool for studying DNA modifications.

## Introduction

Genomic DNA is under constant assault from various damaging agents, leading to breaks and chemical modifications such as oxidation. Among the oxidized base adducts that have been identified [1, 2], 8-oxo-7,8-dihydro-2’-deoxyguanosine (8-oxo-dG), is the most abundant, since guanine, out of the four bases, has the lowest redox potential [3]. Oxidation of G to 8-oxo-dG can occur both directly in the DNA or in the free nucleotide pool by several processes, including the formation of hydroxyl radicals derived from endogenous reactive oxygen species [4], as well as exogenous sources like ionizing radiation [5], and incorporated into the DNA by several polymerases [6]. Its most pivotal characteristic is its ability to both pair with cytosine (forming a regular Watson-Crick base pair) and adenine (forming a Hoogsteen base pair). 8-oxo-dG, when paired with cytosine, is proactively excised from the DNA in humans by the DNA glycosylases OGG1 [7], and adenine, when paired with 8-oxo-dG, is excised by MUTYH [8], but also via preemptive sanitisation of the nucleotide pool by MTH1 (also known as NUDT1) [9]. However, upon failure to repair a 8-oxo-dG:C pair prior to replication, it can lead to a 8-oxo-dG:A pair, which upon a second round of replication would become T:A, leading to a C>A transversion [10, 11]If 8-oxo-dG is incorporated from the nucleotide pool opposite of adenine, then a T>G transversion can also occur after replication [12].Mutations downstream of 8-oxo-dG have significant implications in the development of cancer [13, 14, 15]. 8-oxo-dG is the proposed mechanism underlying COSMIC [16] signatures 18 and 36 [17, 18, 19], and has been recognized as a potential disease biomarker in the field [20, 21]. 8-oxo-dG has also been linked to transcriptional and epigenetic regulation [22], in particular in the context of DNA (de)methylation at cytosine (5-methylcytosine, 5-mC): 8-oxo-dG passively inhibits methyltransferases [23], and OGG1 recruits TET enzymes which convert 5-mC to 5-hydroxymethylcytosine (5-hmC) as the first step in the demethylation process [24].

Due to its potential as a disease biomarker, several efforts have been made to quantify 8-oxo-dG in urine [25, 26], blood [27], genomic material [28], and several other tissues [21]. The absolute quantification of 8-oxo-dG has predominantly relied on highly sensitive methods such as liquid or gas chromatography followed by mass spectrometry or electrochemical detection, or enzyme-linked immunosorbent assays [21]. However, its accuracy has been subject to debate due to large discrepancies between reported levels [29, 30, 31, 21], and high variability between laboratories [32]. Furthermore, these methods fail to provide insights into the genomic location of 8-oxo-dG, which precludes unveiling the mechanisms underlying heterogeneous mutation rates, repair mechanisms, and the role of 8-oxo-dG in epigenetic regulation. For this reason, recently several innovative genomics-based methods have attempted to investigate 8-oxo-dG in a genome-wide manner. These approaches include the detection of apurinic sites created by 8-oxo-dG repair enzymes (Click-code-seq [33] and OGG1-AP-seq [34]), ChIP-seq techniques employing an 8-oxo-dG antibody (OxiDIP-seq [35]), pull-down of cross-linked biotin tags attached to 8-oxo-dG (OG-seq [36] and CLAPS-seq [37]), and pull-down of 8-oxo-dG repair enzymes (enTRAP-seq [38]). Collectively, these methods have revealed that 8-oxo-dG, and its repair, is not uniformly distributed throughout the genome, although with some contradictory results between the methods [39, 22]. Current genomic approaches have three main downsides: first, they lack single nucleotide resolution (with the exception of Click-code-seq), which hampers the study of the relationship between 8-oxo-dG and its associated mutational signatures; secondly, these methods rely on short-read sequencing methods and therefore cannot properly investigate genomic repetitive regions; and lastly, indirect detection is associated frequently with false positive (FP) signals due to suboptimal antibody specificity, enzymatic or chemical reactivity [39]. The latter is especially important given the reported rarity of 8-oxo-dG (1-100 8-oxo-dG per 1 million G [21]). For example, even with a usually considered low false positive rate (FPR) of 1%, the ratio between false and true positives would be approximately 10:1, which would quickly obfuscate any real signal and preclude meaningful conclusions [40].

Nanopore sequencing operates by threading a DNA (or RNA) molecule through a membrane-embedded pore while measuring fluctuations in the electrical current over the membrane, and is currently commercialized by Oxford Nanopore Technologies (ONT) [41]. Changes in the electrical current are indicative of the molecule’s chemical properties, which holds the potential to detect base modifications. This enabled the detection of 5-mC using α-Hemolysin pores [42], and later using ONT devices [43, 44]. Since then, base modification detection models have been developed both by ONT and the community for detecting both naturally occurring and synthetic modifications [45]. But so far, there is no model available for the detection of 8-oxo-dG by nanopore sequencing. Developing modification detection models presents substantial challenges, especially for rare modifications. Firstly, obtaining sequencing data where the precise location of a rare modification is known is not trivial, as these modifications cannot be verified with other technologies to establish a ground truth. Moreover, the context of the modification must exhibit sufficient diversity to prevent the model from learning sequence biases. The effect of the modification passing through the pore on the electrical current must be pronounced enough to distinguish it from inherent noise and other bases. And finally, the model must be extremely precise to accommodate detection of rare modifications, e.g. while for 5-mC it is acceptable to have a 1% FPR, it would be prohibitively high for rare modifications such as 8-oxo-dG due to its low abundance [40].

Keeping these considerations in mind, we devised an approach to generate a library of long synthetic DNA molecules that each contain 8-oxo-dG in a known, specific, but variable sequence context. Utilizing this ground truth dataset, we demonstrate that 8-oxo-dG has a discernible impact on the nanopore raw signal, leading to systematic errors using the ONT provided basecaller. We trained a deep learning model capable of detecting 8-oxo-dG from the raw signal with single-nucleotide resolution, high specificity, and employed it in a cell line with inducible, localized oxidative stress in the nucleus. Our experiments show genome wide variability in 8-oxo-dG levels, with increased levels in complex and repetitive regions which were uncharted by previous short-read based methods. For the first time, we are able to simultaneously measure 5-mC and 8-oxo-dG at single molecule resolution, revealing a strong 5-mC depletion within a 2-kilobase window surrounding 8-oxo-dG. Collectively, our approach showcases a methodology widely applicable to any synthesizable DNA base modification.

### 8-oxo-dG is detectable using nanopore sequencing

To generate a ground truth dataset, we designed a set of 110 short synthetic oligos (46 base pairs each) that contained 8-oxo-dG in different genomic contexts. The oligo design (**Figure 1a**) consists of three barcodes (7 base pairs each) to enable multiplexing. In addition, these barcodes have been optimized to facilitate signal segmentation. The oligos were designed to contain complementary overhangs (10 base pairs) enabling concatenation via hybridization and ligation to create long reads that can efficiently be sequenced on the nanopore platform (**Methods, oligo design**). The 8-oxo-dG base is surrounded by two predefined bases on either side (K_1_, K_2_, K_3_, K_4_), and five additional random bases (2 bases on the 5’ end (N_1_, N_2_), and 3 bases on the 3’ end (N_3_, N_4_, N_5_)) to ensure context diversity. We decided to use predefined bases next to the 8-oxo-dG base, instead of random bases, because the effect of the modification would make it impossible to accurately determine which true bases corresponded to each oligo (**Figure 1b**). Knowing the true sequence of an oligo is necessary to train a basecaller. Finally the oligos include an unmodified guanine base surrounded by the same bases as the 8-oxo-dG base to serve as a built-in control (**Figure 1a**, **Supplementary table 1**).

**Figure 1:**
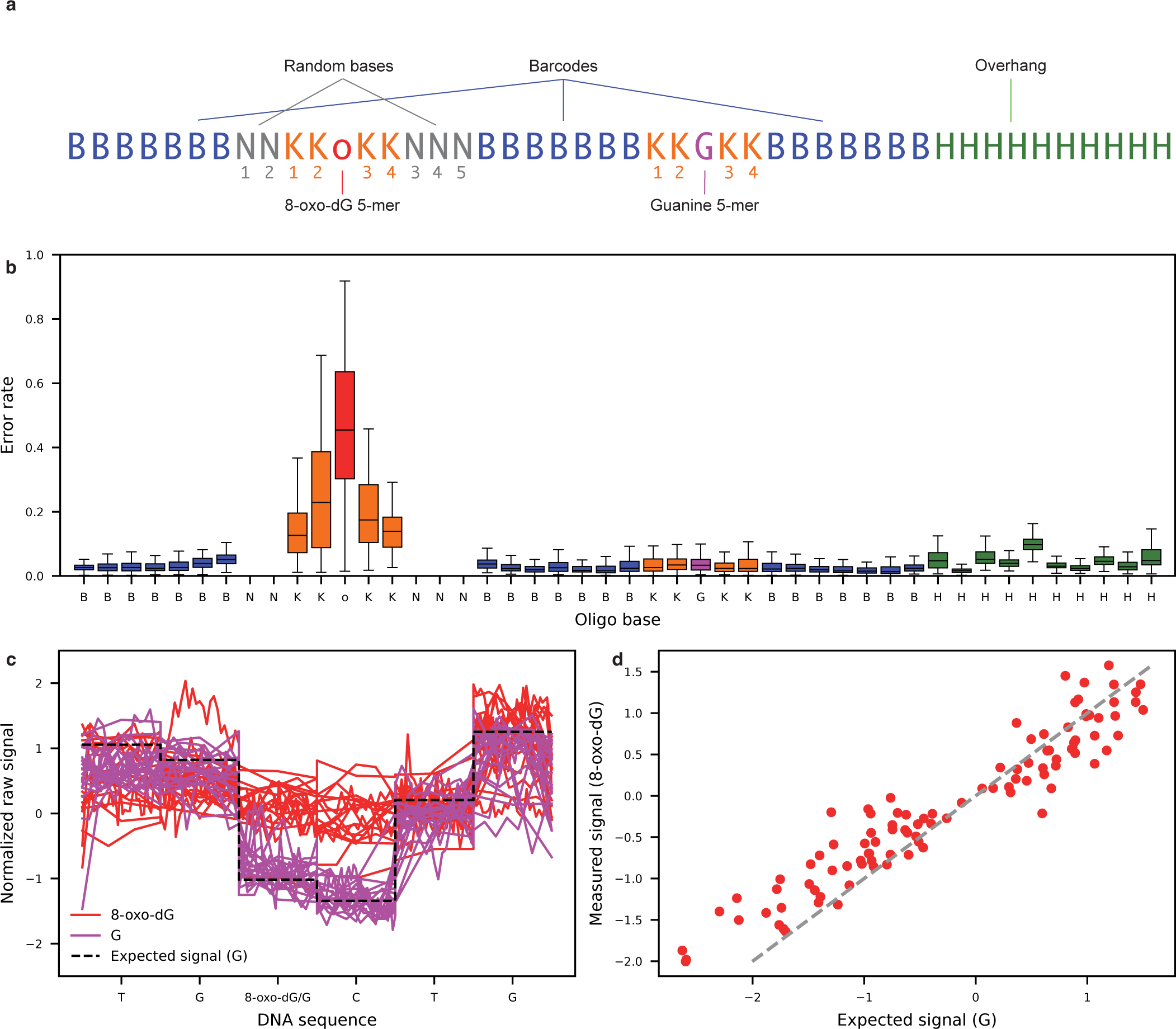
8-oxo-dG has a detectable effect on the nanopore raw signal. **a**, Schematic of the design of the 8-oxo-dG containing oligos. **b**, Error rate per oligo base across all sequenced 8-oxo-dG containing repeats. Random bases are excluded from the analysis as we do not know their true reference. Horizontal bar represents the median, boxplots minimum and maximum bounds represent the 25th and 75th percentiles, respectively, and whiskers extend to 1.5 times the interquartile range. **c**, Example of 8-oxo-dG (red) and G (purple) signal in the TG(8-oxo-dG/G)CTG context. Dashed black line indicates the expected signal value based on the G containing sequence. **d**, Average measured normalized G signal and 8-oxo-dG signal per measured 5-mer as segmented using Tombo. Identity line indicated as the dashed gray line.

Using the concatenated synthetic oligos, we first established whether 8-oxo-dG has an effect on the basecalling performance of ONT’s standard base calling model. We found that there was a substantial increase in basecalling errors specifically at the site of 8-oxo-dG (not basecalling 8-oxo-dG as G) and its neighboring bases (**Figure 1b**). The basecaller’s most likely mistakes, regarding 8-oxo-dG, are deletions (16.4%) or incorrect calls as cytosine (13.5%) or adenine (12.5%) (**Supplementary figure 1**). Moreover, the basecalling error rate is not uniform across different contexts, varying from 1.4% to 91% (**Supplementary table 2**). This variability suggests that the signal alterations are 5-mer context dependent. To establish that the increased error rate was not an artifact of our oligo generated data, we also assessed the error rate on the reverse strand, which is devoid of any modified bases. As expected, here we did observe a consistently low error rate, which was much more uniformly distributed across the different bases and contexts (**Supplementary figure 2**). Interestingly, the cytosine opposite to the 8-oxo-dG base exhibited a slightly higher error rate (median 12.3%) compared to the other bases (median 5.0%) (**Supplementary table 3**). Note that the DNA is unwound when it enters the pore, and hence this cytosine is no longer bound to 8-oxo-dG when it is being sequenced. We hypothesized that the unwinding of the DNA by the helicase might be affected by 8-oxo-dG. We therefore compared the speed (based on raw to expected signal alignment) between cytosines paired to 8-oxo-dG and cytosines paired to G in the same 5-mer contexts. We observed that on average cytosines paired to 8-oxo-dG had fewer measurements, suggesting faster translocation of the DNA, than cytosines paired to G (77% of evaluated 5-mers) (**Supplementary figure 3**). This suggests that, despite only sequencing one DNA strand, the opposite strand can have an impact on translocation speed via structural effects that impact the proteins in the pore, thus affecting base calling accuracy.

We then evaluated what specific effect 8-oxo-dG has on the raw signal. To this end, we aligned the measured raw signal to the expected signal (obtained from ONT’s k-mer model) based on the known underlying sequence (**Methods, Raw data alignment**, **Supplementary note 1**). We observed a significant difference between the expected (which assumes an unmodified guanine) and measured signals. For example, in **Figure 1c** it can be seen that the measured signal is higher than expected when 8-oxo-dG is present. This effect is also clear when evaluating other sequence contexts (**Figure 1d**), indicating that 8-oxo-dG has a clear effect on the measured signal. The measured signal is also significantly different from the other 3 bases (A, C and T) (**Supplementary figure 4**, **Supplementary figure 5**) and is most dissimilar to the pyrimidines. Considering these observations, we concluded that 8-oxo-dG is discernible from the other four canonical bases, suggesting that training a machine learning algorithm for its detection would be feasible.

### *Bonito* basecaller fine-tuning

The nanopore signal originating from 8-oxo-dG containing sequences appears to have several context dependent distinctive features. We therefore sought to train a neural network model that could distinguish 8-oxo-dG from G. We opted for a two step approach, similar to ONT’s *Bonito* + *Remora* combination (**Figure 2a**). First, *Bonito* performs regular basecalling (A, C, G, T); and afterwards, *Remora* classifies the base of interest (e.g. C vs 5-mC, or in our case G vs 8-oxo-dG) as modified or not, using a small data window (e.g. 100 data points).

**Figure 2:**
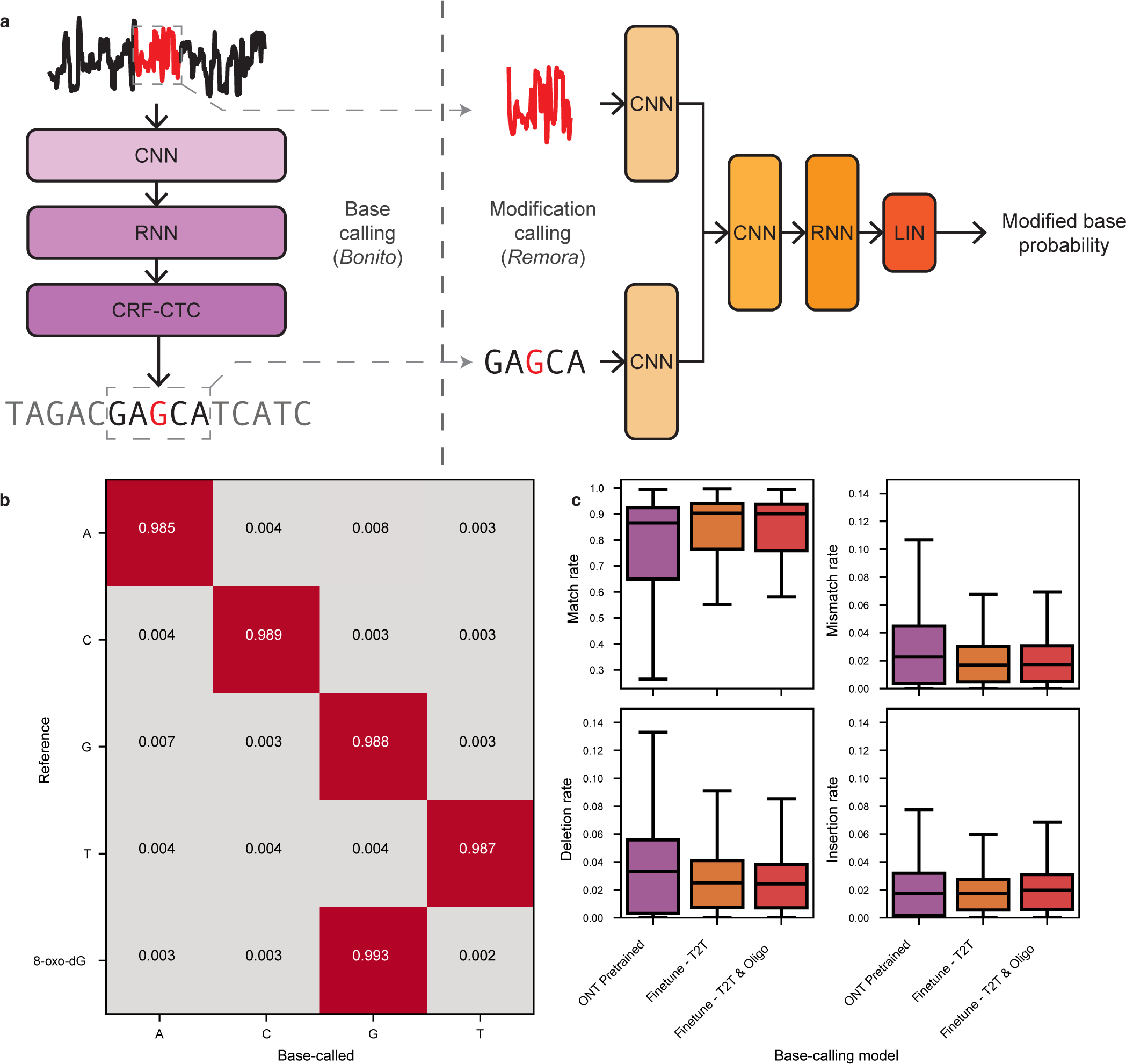
Fine-tuning of a *Bonito* model to basecall 8-oxo-dG. **a**, Schematic of a two step approach to call modified bases in nanopore sequencing. First, the raw signal is basecalled to a DNA sequence. Upon basecalling the (potentially modified) base of interest, a small window of raw signal that corresponds to that particular base is cut; and together with a portion of the basecalled sequence, is given as input to a second model that predicts whether the base is modified or not. **b**, Confusion matrix of the fine-tuned *Bonito* base caller. Values indicate the fraction of outcomes for each ground truth base. **c**, Match, mismatch, deletion and insertion rates of the pre-trained and different fine-tuned models using different datasets on the T2T human reference genome nanopore data. Horizontal bar represents the median, boxplots minimum and maximum bounds represent the 25th and 75th percentiles, respectively, and whiskers extend to 1.5 times the interquartile range.

Since we already observed that the basecaller provided by ONT is not 8-oxo-dG aware, (**Figure 2b**), we first fine-tuned a *Bonito* model to basecall 8-oxo-dG as G. To that end, we used a publicly available pre-trained version of *Bonito* from ONT, and fine-tuned it by training it with oligo concatemers containing 8-oxo-dG and publicly available human data from the telomere-to-telomere (T2T) reference dataset (**Methods, Genomic DNA: Telomere-to-telomere**). We also fine-tuned a model with only data from the T2T dataset to assess the effect of fine-tuning itself on basecalling performance. We then evaluated the basecalling performance of these models on human genomic data, and their capacity to basecall 8-oxo-dG as G, in a cross-validated manner (**Methods, Base calling cross-validation**). We observed that after fine-tuning the models with oligo data, 8-oxo-dG was basecalled as G (*∼*3% median error rate), and immediate neighboring bases are basecalled correctly at error rates similar to the rest of the bases (**Figure 2b**, **Supplementary figure 6**). To ensure that the model did not overfit to the oligo data, we also evaluated the basecalling performance on human genomic data. We observed that basecalling was slightly more accurate on the fine-tuned model than on the pre-trained model (median error rate decreased from 14% to 10%); and that the addition of oligo data in the training did not have a negative effect on regular genomic basecalling (**Figure 2c**). Our *Bonito* fine-tuned model can now accurately basecall 8-oxo-dG as G, and is also capable of regular genomic basecalling.

### Detecting 8-oxo-dG with high specificity

We then trained a *Remora*-like model to distinguish 8-oxo-dG from unmodified guanine. We started by training a base model which takes as input both 100 data points of signal and the basecalled 7-mer. Both features were centered around the guanine of interest. The model then outputs a score between zero and one for the likeliness of that particular guanine being 8-oxo-dG (**Supplementary figure 7**). We trained the model on both positive and negative examples of 8-oxo-dG from our synthetic oligo dataset, as well as on T2T data (**Methods, Modification calling cross-validation, Remora training**). This initial model achieved 93.3% accuracy and 94% specificity at the 0.5 score threshold, with an area under the curve (AUC) of 0.97 (**Figure 3a**). This level of performance is comparable to state-of-the-art 5-mC calling [46]. Problematically, 8-oxo-dG is not as prevalent as 5-mC, and requires a model with near perfect specificity to ensure reasonable FDRs (**Supplementary table 4**). For example, assuming 1-100 8-oxo-dG per million guanines [21], the base model (specificity of 98.6% at the 0.9 score threshold), would be making 100-10000 false positive (FP) calls per true positive call, thus obscuring any real signal. Ideally, we would have a specificity of at least an order of magnitude lower than the prevalence of 8-oxo-dG (Q50-Q70, i.e. 1 false positive per 0.1-10 million guanine calls; (**Methods, Specificity Q-Scoring notation**)). We subsequently explored whether candidate filtering based on basecall errors, or metric learning approaches would improve performance, but we did not observe any major improvements (**Supplementary note 2**).

**Figure 3:**
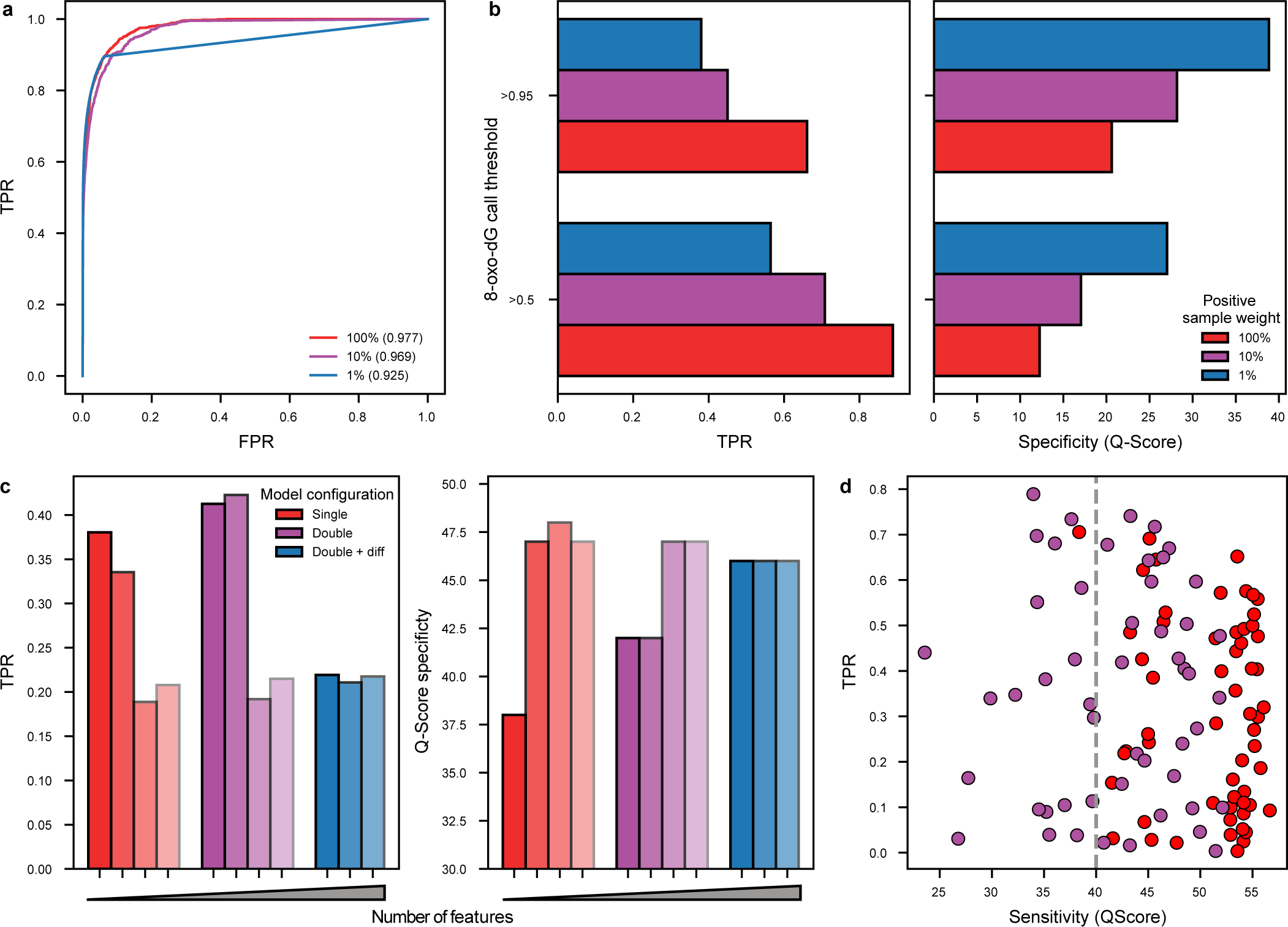
Performance of a *Remora* model. **a**, ROC curves on three Remora models with different positive class weights (100%, 10% and 1%), values indicate the AUC. The straight line shape of the receiver operator characteristic (ROC) curve for the 1% weight model was due to a small number of negative samples with a high 8-oxo-dG score, which reduces granularity in the ROC curve thresholds (**Supplementary figure 8**). **b**, TPR and Q-Score specificity evaluated on the test fold for two different thresholds (0.5, and 0.95) on the three Remora models with different positive class weights (100%, 10% and 1%). **c**, TPR and Q-Score specificity evaluated on the test fold for the experiment in which additional features were added sequentially. Metrics are calculated using a 0.95 threshold. Models include additional features in a cumulative manner, from left to right: basecalls, expected signal, difference between expected and measured signal, and basecall phred quality scores. Red bars include features from the fine-tuned model, purple bars also include features from the Bonito pre-trained model, blue bars include the difference between the features of the Bonito fine-tuned and pre-trained models. **d**, Performance of the Remora model with expected signal as feature per 5-mer at the >0.95 score threshold. Red colored dots indicate 5-mers for which there were no false positive calls, for these 5-mers the QScore was annotated as if a single false positive call was made.

To further improve the specificity of our Remora-like model, we explored the use of class weights during training to emphasize the importance of true negatives (TN). For this purpose, we trained two additional models in which the positive class (8-oxo-dG) had a weight of 10% or 1%, relative to the negative class (G). Using a lower weight on the positive class had a negative impact on global performance, as the AUC was reduced to 0.96 and 0.92, respectively (**Figure 3a**). However, it increased specificity from Q12 to Q27 when using the 0.5 cutoff; and is further increased from Q20 to Q38 when using the more stringent 0.95 cutoff. However, the true positive rates (TPR) decreased from 88% to 56%, and from 66% to only 38% respectively (**Figure 3b**). These results indicate that reducing the weight of the positive class leads to a significant increase in specificity at the cost of an increase of false negatives.

We finally explored if additional signal and sequence features could improve model performance. For example, a signal feature would be the expected signal given the basecalled sequence, and a sequence feature would be the base calling quality scores (**Methods Remora feature-engineering**). We evaluated these features in a feature expansion experiment in which we sequentially added one additional feature at a time. We trained these models with a 1% weight on positive samples since our objective was to further reduce the FPR. Our results indicate that, after adding the expected signal as a feature, the specificity increased from Q38 to Q47 with only a mild 5% reduction of the TPR. Adding additional features, such as the difference between measured and expected signal, and the phred quality scores further increased specificity (to Q48) but with a TPR reduction from 40% to 20% (**Figure 3c, red**). We then added the same features, but based on the pre-trained Bonito base calling model, since its base calls would differ mostly if an 8-oxo-dG was involved. By adding the pre-trained model basecalls and expected signal, the TPR increased by 3%, but did not achieve any specificity improvements (**Figure 3c, purple**). We finally added the difference between the features of the fine-tuned and pre-trained models. These models had similar performance (specificity Q47) and the TPR was at similar levels as the previous models (21%) (**Figure 3c, blue**). Models which contained the expected signal achieved significantly worse TPRs, and we observed that these models greatly overfit during training (**Supplementary figure 9**). We hypothesize that some of these features are predictive but easy to overfit to. For example, just by using the difference of expected signal based on the basecalls from the two models, one can achieve an AUC of 0.79 (**Supplementary figure 10**). In conclusion, the largest improvement was derived by adding the expected signal based on the basecalled sequence as it increased specificity the most without compromising the TPR too much, which also holds true for the less stringent threshold of 0.5 (**Supplementary figure 11**).

Our models have demonstrated an FPR within the same order of magnitude as the expected 8-oxo-dG levels. However, we hypothesized that this error may not be distributed uniformly across the different k-mers, and some k-mers might have an increased FPR. We decided to use k=5 since it is the maximum number of bases with a known reference around 8-oxo-dG in our oligo data. Indeed, four 5-mers demonstrated an AUC of approximately 0.5, equivalent to random guessing. In contrast, 73 other 5-mers exhibited an AUC exceeding 0.9, with the remaining 5-mers falling in between (**Supplementary figure 12**). We also observed that there is a large spread in terms of specificity, ranging from Q24 to Q56 (median Q47) (**Figure 3d**). Notably, for some 5-mers (58 out of 110, indicated in red in **Figure 3d**), we did not detect any false positives, and we calculated their specificity as a single false positive pseudocount. We also observe that the performance spread is also large in terms of sensitivity (average 0.33 *±* 0.22 s.d.), meaning that on average, one out of three 8-oxo-dG molecules will be detected. Because of the low abundance of 8-oxo-dG, we decided to only consider 5-mers that reached a minimum specificity of Q40 (87 out of 110) for subsequent analysis to ensure that the signal to noise ratio is maximized, which corresponds to *∼*35% of the guanines in the human genome.

### H_2_O_2_ production close to the DNA increases 8-oxo-dG levels

Using our nanopore based 8-oxo-dG modification caller, we sought to explore 8-oxo-dG’s locations genome-wide, including repetitive regions previously unexplored by short-read sequencing techniques. To this end we used hTert-immortalized RPE1 cells that express D-amino acid oxidase (DAAO) from *R. gracilis* as a fusion with Histone H2B (RPE1-hTERT-DAAO^H2B^). Upon administration of D-Alanine (D-Ala), DAAO produces H_2_O_2_ in the vicinity of DNA, given its fusion to H2B. Exposure to D-Ala has been shown to give rise to C>A mutations in this mode (in a p53 KO background), suggesting that DAAO^H2B^ activation leads to formation of 8-oxo-dG in the genome [47]. RPE1-hTERT-DAAO^H2B^ cells were exposed to a 2 hour pulse of 20 mM D-Ala, after which we harvested and sequenced the DNA and analyzed it using our 8-oxo-dG model (**Methods, Cell line sequencing**).

We observed that H_2_O_2_ production at the chromatin resulted, somewhat unexpectedly, only in a modest increase of 12 additional detected 8-oxo-dG modifications per a million guanines (16% increase, p-value=0.016) as compared to control (**Figure 4a**). We noted that RPE1-hTERT-DAAO^H2B^ p53^-/-^ cells had overall higher levels of 8-oxo-dG already in control treated cells, compared to RPE1-hTERT-DAAO^H2B^ p53 wild-type, suggesting that the loss of p53 leads to general higher levels of 8-oxo-dG, either by enhanced oxidative stress or slower removal of the modification. After induction of H_2_O_2_ production, both p53^-/-^ and wild-type cells reach similar total 8-oxo-dG levels (**Supplementary figure 13**). We correlated 8-oxo-dG levels to GC content in 1kb bins (**Figure 4b**). We observed that GC content was highly correlated with 8-oxo-dG levels in all conditions (**Supplementary figure 14**). Although this might be expected, we noticed that the rate between 8-oxo-dG and GC content changed at approximately 50%. This indicates that high GC content regions get more easily oxidized, or less effectively repaired, based on the rates measured at lower GC content regions (**Supplementary figure 15**). We next evaluated whether 8-oxo-dG levels were depleted or enriched in specific regions in the genome (**Figure 4c**). Overall, we observed differences in 8-oxo-dG levels depending on the DNA strand (**Supplementary figure 16**) and the chromosomal arm (**Supplementary figure 17**). The forward strands of chromosomes 5, 18 and 19 (p-arms) and chromosomes 1 and 19 (q-arms) have a significant (p-value*<*0.05) 14% or more 8-oxo-dG compared to their reverse strand counterparts, and the reverse strands of chromosomes 12, 14, 15 (p-arms) and chromosome 16 (q-arm) have 9% or more 8-oxo-dG compared to their forward strand counterpart. These particular differences are treatment and p53 status independent (**Supplementary figure 18, Supplementary figure 19, Supplementary figure 20, Supplementary figure 21**). We then analyzed the relative 8-oxo-dG levels across different genomic regions (**Figure 4d**). Again, we observed a global increase in 8-oxo-dG levels after H_2_O_2_ production, irrespective of genomic region (**Supplementary figure 22**). Non-repetitive and centromeric genomic regions showed 8-oxo-dG levels that roughly align with the overall rate of *∼*80 8-oxo-dG per 1 million G (**Figure 4d, gray dashed line**); with the exception of the 5’-UTR, which has the highest median levels, as also previously reported by Ding et al [36]. On the other hand, complex repetitive and satellite regions contained, relatively, the most 8-oxo-dG with high variance between chromosomes (**Figure 4d**). We fitted a linear model to evaluate whether 5-mer content, or the relative abundance of the different genomic regions, were causative for these observations, however none could explain the observed variability (**Supplementary note 3**).

**Figure 4:**
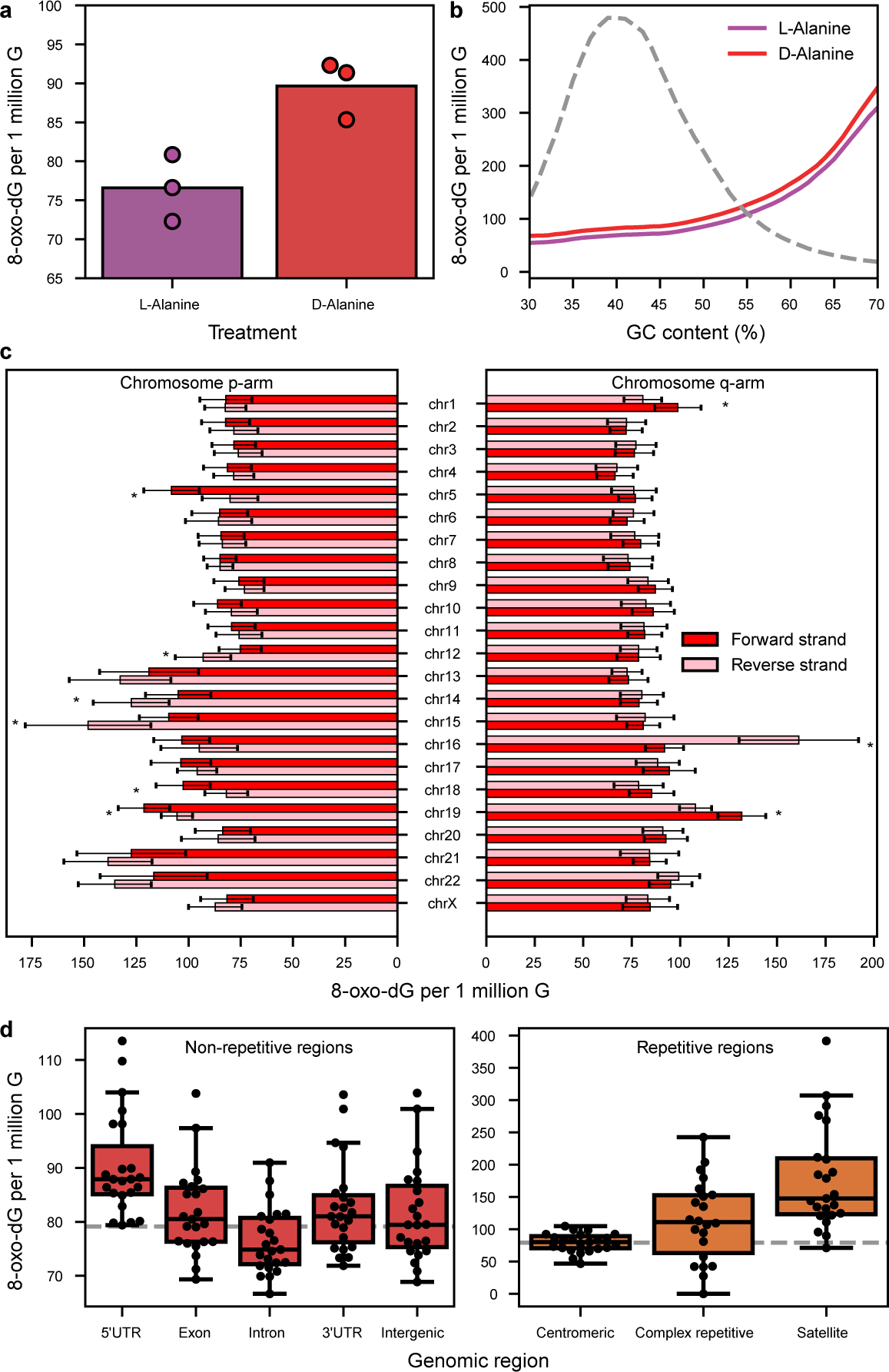
8-oxo-dG distribution across the genome. **a**, Overall 8-oxo-dG molecules per 1 million G molecules per L-Alanine and D-Alanine treated cells. Error bars indicate minimum and maximum calculated values. Dots indicate the values per sequenced condition. **b**, 8-oxo-dG levels across different GC (%) content bins. Blue and red lines indicate values for L-Alanine and D-Alanine treated cells respectively. Grey dashed line indicates the distribution of measured GC content bins. **c**, 8-oxo-dG levels per chromosome, chromosome arm and DNA strand. Bars indicate average values. Error bars indicate the minimum and maximum calculated values across all conditions. Asterisks indicate a significant p-value (*<*0.05) derived from a two-sided t-test between the values of the forward and reverse strands. **d**, Distribution of 8-oxo-dG levels per genomic region type across all conditions and chromosomes. The dashed gray horizontal line indicates the overall 8-oxo-dG level across all conditions, irrespective of genomic region. Horizontal bar represents the median, boxplots minimum and maximum bounds represent the 25th and 75th percentiles, respectively, and whiskers extend to 1.5 times the interquartile range. Black dots indicate the underlying data.

Finally, we evaluated 8-oxo-dG at telomeric regions since they are guanine rich, and their oxidation has been linked to senescence and telomere shortening [48, 49]. To that end, we combined the sequencing data from all experimental conditions, and obtained a total of 487 reads that primarily mapped to the telomeres. We observed a strong bias towards reads mapping on the p-arm reverse strand and q-arm forward strand (**Supplementary figure 23**); which can be explained by blunt end requirement of the ligation mechanism in the library preparation (**Supplementary note 4**). Notably, we only detected 2 high confidence 8-oxo-dG calls on the q-arm telomeres of chromosomes 16 and 19 (**Supplementary figure 24**), a value ten times lower than expected, given the 50% GC content of the region. This result might indicate that 8-oxo-dG is very efficiently repaired at telomeres as a preventive measure to downstream complications.

### The 8-oxo-dG trinucleotide context profile does not match the C>A mutational signatures

Because of its ability to mispair with A, unrepaired 8-oxo-dG present at the time of DNA replication, oxidative stress has been the proposed mechanism behind the COSMIC mutational signatures 18 and 36 [17, 18, 19]. We therefore wanted to compare C:G>A:T (here mentioned solely as C>A) mutations to the derived 8-oxo-dG profile to establish if the resulting mutations had the same trinucleotide context as the detected 8-oxo-dG. Using the same RPE1-hTERT-DAAO^H2B^-p53^-/-^ cell lines, van Soest et al. [47] performed Illumina sequencing to analyze the mutational profile caused by H2B-DAAO-derived H_2_O_2_. This analysis was also performed in a p53 WT background in a parallel experiment not shown in van Soest et al. [47]. We therefore used these mutational profiles and compared them to the 8-oxo-dG profile. It must be noted that the mutational profiles were obtained after 4 rounds of 20mM D-Ala treatment and recovery; while for our 8-oxo-dG analysis, we did a single 20mM D-Ala pulse followed by immediate DNA harvesting. We therefore abstain from drawing any strong conclusions regarding the relationship between number of mutations and 8-oxo-dG due to the treatment difference.

The mutational profiles were highly similar in terms of COSMIC mutational signatures present (minimum cosine similarity of 0.95) (**Supplementary figure 25**), irrespective of H_2_O_2_ induction and p53 status. The only major difference between the samples was the amount of mutations [47]. Since our aim is to compare the mutational signatures, we combined the mutational profiles of all the sequenced conditions (**Figure 5a**). To make the mutational profile comparable to the 8-oxo-dG profile, we recalculated the mutational profile including only mutations in the 5-mer contexts in which our model performed with at least Q40 specificity, which reduced the total number of C>A variants from 6880 to 2506. We also normalized the relative contribution of each 3-mer based on the number of 5-mers that passed the specificity threshold (**Supplementary figure 26**). We observed that these limitations did not drastically change the originally derived mutational profile (**Supplementary figure 27**).

**Figure 5:**
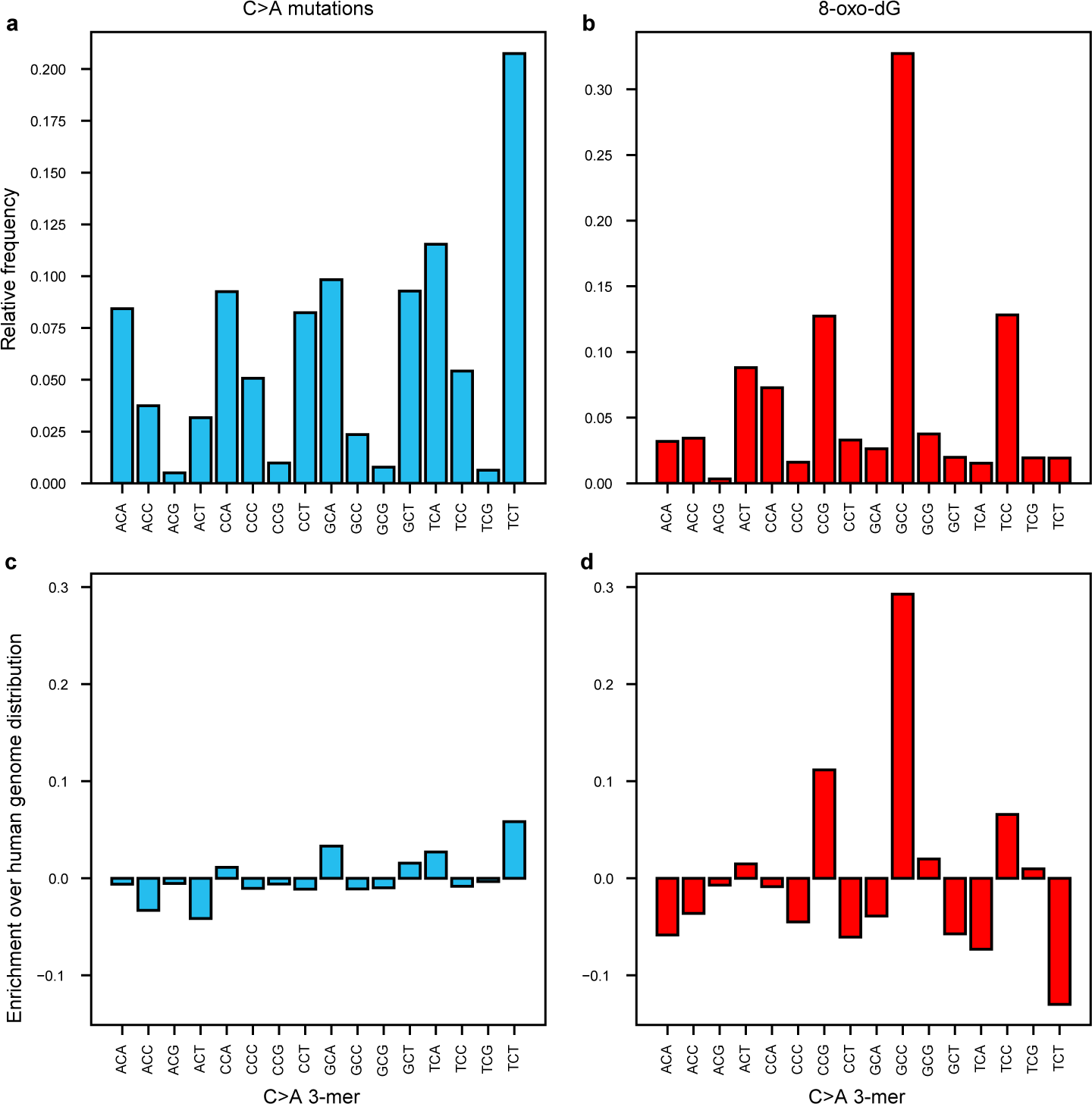
Mutational and 8-oxo-dG signatures. **a**, Combined mutational signature (C>A or G>T) from all the cell lines derived from Illumina sequencing. **b**, 8-oxo-dG normalized abundance profile for each 3-mer. Note that the 3-mers are annotated as the reverse opposite strand of 8-oxo-dG (e.g. ACT would be equivalent to AXT, where X denotes 8-oxo-dG). **c**, Mutation enrichment of each 3-mer normalized to the abundance of each 3-mer in the human genome. **d**, 8-oxo-dG enrichment of each 3-mer normalized to the abundance of each 3-mer in the human genome.

We detected many more 8-oxo-dG bases than C>A mutations in every analyzed 3-mer context despite the shorter treatment (**Supplementary figure 28**). The C>A mutational profile (**Figure 5a**) and the 8-oxo-dG profile (**Figure 5b**) however only moderately agree (cosine similarity of 0.33). Our results do therefore not discard the hypothesis that 8-oxo-dG is the underlying cause of C>A mutations, but rather that the rate at which 8-oxo-dG leads to a C>A mutation is 3-mer context specific. Notably, the C>A mutational profile has a high similarity (cosine similarity of 0.95) with the 3-mer abundance profile of the human genome (**Supplementary figure 29**) and the observed mutation profile thus seems to more closely relate to 3-mer abundance than to 8-oxo-dG location itself (**Figure 5c**). This suggests that C>A mutations are likely driven by other or additional mechanisms such as replication timing [50]. Interestingly, we do observe strong 8-oxo-dG enrichment and depletion for certain 3-mers (**Figure 5d**), in particular 3-mers that contain a CG or a GG motif. This result coincides with our previous observation of 8-oxo-dG enrichment in high GC content regions, but might hint at other mechanisms such as a relationship with CpG methylation.

### 8-oxo-dG levels negatively correlate with methylation levels

Previous work has linked 8-oxo-dG with both inhibition of DNA methylation [23, 51], as well as active demethylation via TET enzyme recruitment by OGG1 [52, 24]. Whereas previous methods for genomic 8-oxo-dG detection precluded the simultaneous assessment of 8-oxo-dG and other base modifications, our approach can, for the first time, look at 8-oxo-dG, 5-mC and 5-hmC on the same DNA molecule. We therefore compared the methylation and hydroxymethylation status of the surrounding CpG sites between G and 8-oxo-dG containing regions in a 10 kilo-base window in the same DNA molecule.

We observed that there is a significant general decrease in methylation levels around oxidized guanines compared to non-oxidized (**Figure 6a**, **Methods, 8-oxo-dG related methylation analysis**). The decrease in methylation levels correlates with distance to the oxidized base: it is lowest close to 8-oxo-dG (*∼*45% methylation), and recovers to average genome-wide methylation levels at approximately 2000 base from 8-oxo-dG (*∼*54% methylation). This effect was observed irrespective of p53 status and DAAO^H2B^-derived H_2_O_2_ production, suggesting a DNA damage independent relationship (**Supplementary figure 30**). We then grouped the data by 3-mer analogous to the profile analysis and observed that the decrease in methylation levels correlated with the 3-mer enrichment for 8-oxo-dG (**Figure 6b-c**). The decrease in methylation is largest for 3-mers with a CG motif, with the exception of CGT. A decrease in methylation is also observed for 3-mers without a CG motif, this is most apparent for GGA, which has a similar decrease in methylation level as compared to CG containing 3-mers. Interestingly, in 3-mers without a CG motif the methylation levels reached average genome wide levels at approximately 1 thousand base-pairs from 8-oxo-dG, while methylation levels in 3-mers with a CG motif did not return to baseline within 5kb from 8-oxo-dG [Figure 6c]. Strangely, the AGG was the only 3-mer that displayed overall higher methylation levels for 8-oxo-dG containing molecules (**Figure 6c**). We observed that 5-hmC levels were slightly higher in 8-oxo-dG vs guanine containing regions (average *∼*3.5% vs *∼*3%) (**Supplementary figure 31**). Similarly to 5-mC, we did not observe p53 or H_2_O_2_ dependent effects (**Supplementary figure 32**). However, upon inspection of the 5-hmC levels per 3-mer, we observed that there is no difference in 5-hmC levels per 3-mer with the exception of two cases (**Supplementary figure 33**). First, 5-hmC levels are highest around the GGA context, reaching an average of 6% in regions containing 8-oxo-dG. And secondly, the opposite effect happens in the CGA context, where 5-hmC levels are highest for regions with G (average *∼*5,8%) (**Supplementary figure 33**).

**Figure 6:**
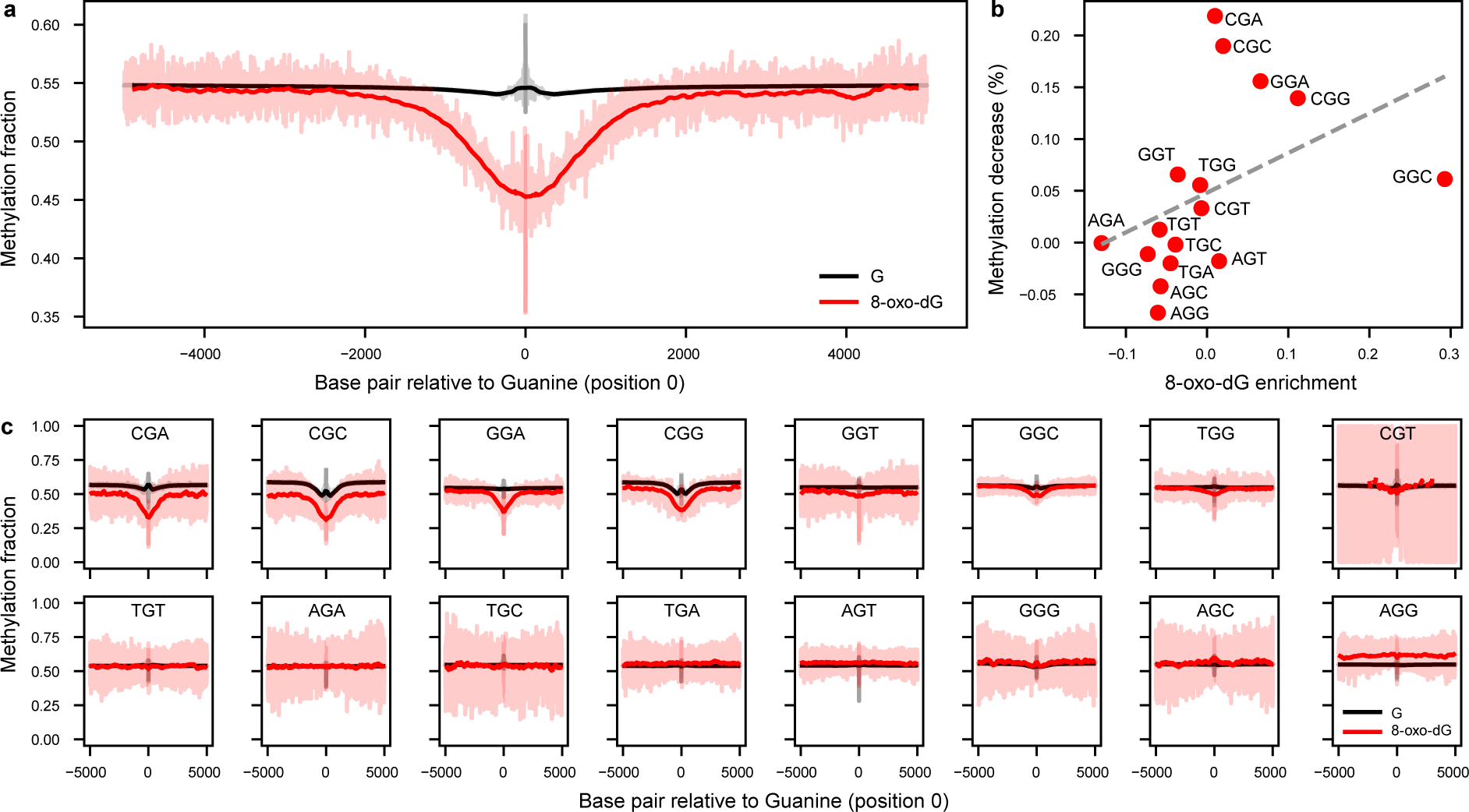
Genomic de-methylation surrounds 8-oxo-dG. **a**, Methylation levels for 8-oxo-dG (red) and Guanine (black) containing reads. Reads are centered around (position zero) the 8-oxo-dG or Guanine. Methylation levels are obtained from the same molecule. The transparent red line indicates the underlying data, the dark gray line is the result of an 11 base average convolution. Data from all experimental conditions is included, see (**Supplementary figure 30**) for the per condition analysis **b**, Correlation between 8-oxo-dG enrichment as in Figure 5d and the methylation difference at position zero. **c**, Similar to panel **a**, but data has been split based on the 3-mer (reverse complement) surrounding the guanine.

Our approach enabled simultaneous assessment of 8-oxo-dG, 5-mC, and 5-hmC on the same DNA molecule for the first time. Together with our analysis of the mutational signatures, our results suggest that 8-oxo-dG abundance has a primary role in epigenetic regulation, and that its mutagenic effect would play a secondary role.

## Discussion

Nanopore sequencing holds the potential to detect any base modification, both on DNA and RNA. However, due to technical challenges, accurate models have only been established for a limited number of modifications [45]. Available models focus on highly prevalent modifications, like 5-mC or 6mA because they can be generated enzymatically, are highly stable, and therefore training data can be easily obtained and verified through alternative sequencing technologies [45]. However, these approaches are not widely applicable to less abundant modified bases [53]. In this study, we leverage synthetic DNA to bridge this gap, enabling the generation of a fully controlled ground truth dataset. We show that our approach can successfully be used to detect 8-oxo-dG, and has the potential to be expanded to nanopore-based detection of other base modifications. Furthermore, compared to existing 8-oxo-dG detection methods, our approach does not require any complex sample preparation, chemical reactions or antibody pull-downs, and can be readily applied to existing nanopore data.

Our work showcases that developing deep-learning models required for detecting rare modifications from the raw nanopore squiggles is not trivial. False positives drastically decrease the signal-to-noise ratio and, if not low enough, will mask any inherent biological signal [40]. We address this by using label weighting and feature engineering. However, while we achieved a major increase in specificity, this comes at the price of reduced sensitivity of the model. Rare base modifications also pose a challenge in regular base calling. As we have shown, 8-oxo-dG naive basecallers have a significantly increased error rate on molecules that contain an 8-oxo-dG modification. Therefore, until these rare modifications are integrated in the basecaller training pipelines, perfect accuracy for all natively sequenced molecules will not be achievable.

Using our highly specific model, we show differential 8-oxo-dG levels across the different chromosomes and genomic regions of RPE1-hTERT cells. Our observations fit with the variability shown in OGG1 driven apurinic sites [34]; and we also observe a similar distribution of 8-oxo-dG over non-repetitive regions as reported in Ding et al. [36]. For the first time, we can evaluate 8-oxo-dG abundance on highly repetitive sequences, which show high levels of 8-oxo-dG, indicating either increased susceptibility to oxidation or less efficient repair. Surprisingly, we observed very low 8-oxo-dG levels at the telomeres despite their oxidation susceptibility [54]. This might suggest that telomeric 8-oxo-dG is very efficiently repaired to avoid telomere shortening and the downstream induction of cellular senescence [48, 49]. However, our coverage was limited, and further research using telomere enrichment techniques [55] might advance our understanding of the repair mechanisms of 8-oxo-dG at telomeres [56].

The mutational process behind C>A mutations in COSMIC signatures 18 and 36 has been long attributed to oxidative stress and unrepaired 8-oxo-dG [17, 18, 19]. Our results show that, primarily, 8-oxo-dG does not strongly correlate with the same nucleotide context of the signatures. Rather, these signatures strongly follow the human 3-mer genomic content, which indicate that the mutational rate of 8-oxo-dG is 3-mer specific, and could be driven by other mechanisms such as replication timing [50]. Importantly, our results do not discard 8-oxo-dG as the cause of C>A mutations, since the modified base is far more abundant than the resulting mutations. Rather, we envision that these mutations are secondary to the role of 8-oxo-dG in epigenetic regulation.

We observed a distinct decrease in CpG methylation levels around 8-oxo-dG, however it is unclear whether CpG demethylation precedes guanine oxidation, or vice-versa. It has been shown that 8-oxo-dG inhibits methyl-transferase enzymes [23, 51, 52], which would indicate that oxidation happens in already de-methylated regions. Another possibility would be that active demethylation is promoted via the recruitment of TET enzymes by OGG1 [24]. At first sight, this model suggests that, 8-oxo-dG then would have been removed by OGG1, and therefore would not be detected in the first place. However, OGG1 oxidation can block its glycosylase and lyase activity, but not its binding to 8-oxo-dG [57], which would explain why we still measure 8-oxo-dG, and indicate that oxidation happens before de-methylation. Finally, some histone demethylases are known to produce H_2_O_2_ as part of the histone demethylation reaction (e.g. H3K9 [58] and H3K4 [59], which could cause the formation of multiple 8-oxo-dGs while oxidatively inhibiting the recruited OGG1. This model would link histone and DNA epigenetics through redox regulation. Perturbation experiments on OGG1, in combination with TET and histone demethylases, would provide valuable insights to further elucidate the underlying regulatory mechanism.

In conclusion, our work showcases a viable methodology for rare modification detection using nanopore sequencing, which could be applicable to detect any synthesizable base modification. These models can then be later used to decipher and further understand epigenetic regulatory mechanisms and the interplay between DNA modifications, as well as potential implications for disease mechanisms and biomarker detection.

## Methods

### Oligo design

A total of 110 oligos were designed in complementary pairs, wherein each forward oligo contains an 8-oxo-dG base and was paired with its complementary reverse strand oligo devoid of any base modifications. Oligos are 46 base pairs long and contain three barcodes, two 8-oxo-dG/G k-mer regions of 10 and 5 bases respectively, and a 10 base overhang. The barcodes are 7 bases long and were defined to have the lowest basecalling error rate possible (based on the T2T dataset (**Methods, Genomic DNA: Telomere-to-telomere)**) and a high sequence entropy (meaning that the same base was never repeated sequentially). The 10 base overhangs are complementary between the forward and reverse strands, which allow concatenation via hybridization and ligation to create long molecules that are more readily sequenced on the nanopore platform. Between the first and second barcode lies an 8-oxo-dG that is immediately surrounded by two known bases at either side, which provides the known sequence context. Similarly, between the second and third barcode a guanine is surrounded by the same four bases. To maximize sequence variability around 8-oxo-dG, five additional random bases were added: two on the 5’ end, and three on the 3’ end.

### Oligo concatenation

Oligos were first annealed in a thermocycler by mixing the two complementary oligos in equimolar rates. Oligos were diluted in 1X T4 ligase buffer (NEB REF #B0202S) in a total volume of 10µL. The mixture was heated to 95°C for 5 minutes, and then the temperature was decreased by 0.1°C per second until 4°C. After hybridization, oligos were concatenated by ligation. 9µL of 1X T4 ligase buffer (NEB REF #B0202S) and 1 µL of T4 ligase (NEB REF #M0202S) were added to the solution. Afterwards, the mixture was incubated at 16°C for 18h. T4 ligase was then inactivated at 65°C for 10 min. The resulting DNA was cleaned using the Qiagen PCR & Gel Cleanup Kit (Qiagen REF #28506) according to the manufacturer’s instructions, and eluted from the column using Milli-Q water. This process was repeated 3 times to increase the concatemers length (**Supplementary figure 34**). Oligo concatemer concentrations were then quantified via Nanodrop (Thermo Scientific NanoDrop 2000 #ND-2000) and multiplexed prior to library preparation in equimolar rates as indicated in **Supplementary table 1**.

### Genomic DNA: Telomere-to-telomere

We used existing nanopore sequencing data from the reference genome (NA12878/GM12878, Ceph/Utah pedigree) dataset [60]. This human dataset contains many different sequencing runs. We arbitrarily chose three experiments so that each different ligation kit (rapid, ligation and ultra) would be included: FAB42828, FAF09968 and FAF04090. We assumed all the sequenced DNA did not contain any 8-oxo-dG as it had been prepared using the standard library preparation protocol, which contains a repair step with Fpg, a DNA glycosylase that removes 8-oxo-dG.

### Library preparation

Oligo concatemers and genomic DNA samples were library prepared using the SQK-LSK109 ligation kit according to the manufacturer’s instructions, with the exception that the FFPE repair step was skipped. The exclusion of this step was deliberate and meant to preserve 8-oxo-dG in our samples since the enzymatic function of Fpg as a DNA glycosylase is responsible for removing 8-oxo-dG from DNA.

### Nanopore sequencing

Samples were sequenced using MinION R9.4.1 flow cells for 72h using the GridION device. MinKNOW v22.12.5 or earlier was used to ensure that all our samples were sequenced using 4KHz sampling. Oligo concatemers were multiplexed as indicated in **Supplementary table 1**. Each genomic DNA sample was sequenced individually in a single MinION flow cell.

### De-multiplexing and reference assignment

Although oligos were concatenated separately, the library preparation protocol contains a ligation step in which all oligo concatemers are pooled together. Therefore, it is possible to get oligo hybrids that contain different reference sequences. For this reason, oligo samples were de-multiplexed using a custom algorithm to detect the individual oligo repeats and barcodes within a read. Given the basecalls of a read, all possible oligo reference sequences within that batch are aligned to the basecalled sequence using a semi-global alignment algorithm (as implemented in the parasail library [61]). The reference sequence with the highest number of matches is considered as the true underlying sequence for that portion of the basecalls. However, only matches to the barcode and guanine 5-mer portions of the reference sequence are considered (max of 26 matches), since the overhang sequence is common for all oligos and the basecalls surrounding 8-oxo-dG cannot be trusted. Then, the aligned portion of the basecalled sequence is masked to avoid further alignment in subsequent iterations. This process is repeated until there are fewer than 15 non-masked bases or more than a set maximum number of iterations is exceeded, which is dependent on the length of the basecalled sequence.

### Raw data alignment

Raw data was aligned to the expected signal based on the reference sequence of the read. We used a custom script that featured Tombo’s API [62] to align the two sequences. Notably, the expected signal model is based solely on non-modified bases and therefore, we expected the raw signal corresponding to the 8-oxo-dG and surrounding bases to contain mis-alignments. Because our oligos contain random bases, for which we do not know their true reference, we used their aligned basecalls instead.

### Base calling cross-validation

The oligo concatemers dataset was first split into train and test sets. The test set consisted of 100 reads (chosen at random) per different oligo sequence (total of 22000 reads). To assess model performance during training, a portion of the train set was split into the validation set at random, and kept constant throughout the epochs (10% of the total training examples). The human dataset was first split into train and test sets. The train set consisted of 40000 reads, chosen at random, from odd numbered chromosomes. The test set consisted of 10000 reads, chosen at random, from even numbered chromosomes. In the same manner as the oligo data, 10% of the total training examples was used as a validation set to assess model performance during training.

### Raw signal normalization

A common approach to normalize the nanopore raw signal is to center and scale its values based on the set of measurements from the whole read (**Equation 1**).

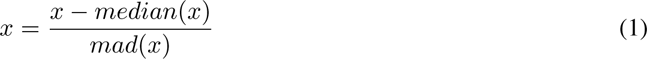

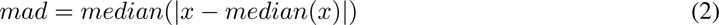

We observed that both the T2T and oligo data had a consistent shift between expected and measured values (**Supplementary figure 35,Supplementary figure 36**). We attributed this larger shift in the oligo data to the consistently repetitive nature of the oligo sequences, which overall, do not have enough sequence variability to guarantee that the standard normalization approach would work as expected. To avoid normalization bias, which could already distinguish genomic from oligo data at the signal level, we performed a second normalization step. In this second normalization, we calculate what would be the optimal median and mad (**Equation 2**) values to minimize the distance between measured and expected values after alignment. We do this by first fitting a linear model using least squares regression between the aligned measured and expected values. We then re-scale the initially calculated med and mad values based on the fitted model. Using this second normalization step, we noted that the previously observed signal shift was corrected, and average 6-mer values match between genomic and oligo data (**Supplementary figure 37**). We applied this 2-step normalization approach on all our data both before training and inference.

### *Bonito* fine-tuning

We fine-tuned a *Bonito* model to basecall 8-oxo-dG as G. We started from a pre-trained open source state provided by ONT in their public bonito repository. We fine-tuned the model for a total of 43000 training steps using a batch size of 256. The model was fine-tuned using an unbalanced combination of oligo (25% of all training examples) and T2T data (75% of all training examples). We used the Adam [63] optimizer with a constant learning rate of 0.0005. Other parameters of the optimizer were: β1 = 0.9, β2 = 0.999, ε = 1e-8 and λ = 0.0. We used a dropout of 0.1 in between CNN layers, and a dropout of 0.5 in between RNN layers (**Supplementary figure 38**).

### Modification calling cross-validation

To avoid any potential data leakage between the fine-tuning of the base calling model and the training of the modification calling model, we used the same (based on read id) train, validation and test data splits in both steps.

### *Remora* training

We trained a *Remora* model to distinguish 8-oxo-dG from a regular G. We used a neural network architecture that first encodes the signal and sequence features via convolution, concatenates the encoded vectors, forwards the concatenated vector through a convolution layer and two recurrent layers, and a final linear layer for classification (**Supplementary figure 7**). We trained several models with balanced (via upsampling of the positive label samples, or downsampling of the negative label samples) and unbalanced datasets. We also trained models with different positive label weights: 100% 10% or 1%, with different features (**Methods, *Remora* feature-engineering**), and a model using metric learning (**Methods, *Remora* metric learning**). All models were trained as described in the following paragraph.

We trained the model as a classical classification problem using cross-entropy as loss function for a total of 235000 training steps using a batch size of 256. We used the Adam [63] optimizer with a variable learning rate, which started at 0.00005 and increased linearly for 5000 training steps until 0.001, and then decreased using a cosine function until 0.00005. Other parameters of the optimizer were: β1 = 0.9, β2 = 0.999, ε = 1e-8 and λ = 0.0. We used a dropout of 0.2 between all layers of the model.

### *Remora* feature engineering

Traditional *Remora* models from ONT include the raw signal and base calls, centered around the base of interest as input features for the model. We explored the use of additional features, both at the signal and base call level. At the signal level, we used the following features: expected signal aligned to the measured signal based on the base calls from the fine-tuned *Bonito* model (8-oxo-dG aware), or from the pre-trained *Bonito* model (not 8-oxo-dG aware); the difference between the measured and expected signals (from both models); the difference between the aforementioned differences. At the sequence level, we used the following features: base calls from both the fine-tuned and pre-trained *Bonito* models, phred quality scores from both the fine-tuned and pre-trained*Bonito* models. To ensure local feature information, features (both signal and sequence level) that were further than 3 bases, on either side, of the target G (based on the basecalls) were masked with zeros.

### *Remora* metric learning

We trained a *Remora* model using Multi-Similarity loss (α=2, β=50, base=0.5) and Multi-Similarity miner (ε=0.1) [64] as implemented in PyTorch-Metric-learning. Triplets provided by the miner were filtered to only contain samples from the same 5-mer in an effort to force the model to compare the two labels in the same sequence context. Cosine similarity was calculated on the embedding vector output of the last LSTM layer (**Supplementary figure 7**) and used as a distance metric to calculate the loss. Triplet loss and cross-entropy loss were added together with equal weights before backpropagation.

### Specificity Q-Scoring notation

Due to the low prevalence of 8-oxo-dG, it is necessary to achieve a near-zero false positive rate (10^-5^-10^-7^). To make the annotation of these very small values easier, we convert these using the phred quality score (Q) [65].

### Cell line sequencing

RPE1-hTERT-DAAO^H2B^ wild type (WT) and p53^-/-^ cells were treated for 2h with 20mM L-Alanine or D-Alanine. Afterwards, genomic DNA was harvested using the DNeasy blood and tissue kit (Qiagen REF #69504) according to the manufacturer’s protocol. Nanopore sequencing libraries were prepared and sequenced in the same manner as described for the oligo concatemers (**Methods, Library preparation, Nanopore sequencing**). For additional information regarding DAAO constructs and Illumina sequencing of these cell lines see van Soest et al [47].

### Illumina derived mutation profile

Clones were sequenced at 30x base coverage using an Illumina Novaseq 6000 or an Illumina Hiseq X10 sequencing machine. Sequencing reads from all samples were mapped to the human reference GRCh38 genome using the Burrows-Wheeler Aligner v0.7.17. Duplicate sequencing reads were marked using Sambamba MarkDup v0.6.8. Variants in the mapped data were called using GATK Haplotypecaller version 4.1.3.0 using default settings. Variants were filtered using GATK 4.1.3.0 using several filter settings **Supplementary table 5**.To filter out mutations induced during sequencing, clonal expansion or library preparation, we filtered genomic variants using an in-house filtering pipeline, SMuRF v2.1.1. Briefly, the variant allele frequency (VAF) was calculated for each variant by pileup of all bases mapped at the mutation position. Variant data derived from cell clones were filtered for the following criteria: VAF *≥*0.3, base coverage *≥*10 and an MQ quality *≥*60. To select only mutations occurring during in-vitro culture, variants present in the clonal parental cell line were removed. Recurrent mapping or sequencing artifacts were removed by filtering against a blacklist containing variants present in healthy bone marrow mesenchymal stromal cells.

### 8-oxo-dG mutational profile analysis

High confidence 8-oxo-dG calls (score > 0.95) from >Q40 5-mers were grouped based on the 3-mer of the opposite strand to facilitate comparison with the C>A profile as described in the COSMIC signatures database. Relative frequencies were calculated as the fraction of counts of each 3-mer given the total amount of counts. Counts per 3-mer were normalized to the amount of training 5-mers per 3-mer. Enrichment was calculated by subtracting the relative frequency of 8-oxo-dG calls from the relative frequency of 3-mers in the T2T reference genome.

### 8-oxo-dG genomic regions analysis

Genomic region annotations were downloaded for the CHM13v2 telomere-to-telomere (T2T) reference genome (https://github.com/marbl/CHM13). Base calls were aligned to the T2T reference genome using minimap2 [66]. 8-oxo-dG counts (score > 0.95) were normalized per coverage as well as per 5-mer relative abundance per region. 8-oxo-dG counts on 5’-UTR, 3’-UTR, exon and intron regions were only considered if the molecule was found on the same DNA strand as the annotated region; in intergenic, satellite, centromeres and complex repetitive regions 8-oxo-dG counts were considered regardless of the DNA strand. 5’UTR was annotated as the upstream 5000 base pairs before the start of all annotated coding sequences. 3’UTR was annotated as the downstream 5000 base pairs of all annotated coding sequences. Intergenic regions were considered as all intervals that had no annotation.

### 8-oxo-dG related methylation analysis

All nanopore sequencing data was methylation called using the guppy tool and ONT’s model *dna*_*r*9.4.1_*e*8.1_*modbases*_5*mc*_5*hmc*_*cg*_*hac.cfg*. Methylation calls were mapped to the T2T reference genome assembly, and then extracted using the modkit tool. Analysis was restricted to only 5-mers in the training set. We evaluated the methylation status within a 10 kilobase window centered around any guanine (canonical or 8-oxo-dG). Methylation prediction scores were then averaged per base pair position to calculate the average methylation status. Methylation scores within a 10 base window around 8-oxo-dG were masked out from the analysis because we assume that the methylation calling model is not aware of signal effects that might be caused by a close proximity 8-oxo-dG molecule.

### 8-oxo-dG telomere analysis

Reads whose mapping was primarily to the annotated T2T telomere regions were used for analysis. Based on the reference T2T genome, annotated telomere regions were further constrained to the last TTAGGG repetitive element.

## Declarations

## Data availability

Nanopore sequencing data for the synthetic oligos and the cell lines has been uploaded to the European Nucleotide Archive under accession code PRJEB76712. Nanopore human data from the Telomere-to-Telomere can be found at: https://github.com/nanopore-wgs-consortium/NA12878/blob/master/Genome.md [60]. Data availability details regarding Illumina sequencing data can be found at van Soest et al [47].

## Code availability

Source code for the 8-oxo-dG caller as a Python package can be found at https://github.com/marcpaga/esox. The following Python v3.7 packages were used during the development of the 8-oxo-dG caller: fast-ctc-decode (v0.3.2), jupyterlab (v3.6.1), mappy (v2.22), matplotlib (v3.5.3), numba (v0.54.1), numpy (v1.18.5), ont-fast5-api (v4.1.1), ont-tombo (v1.5.1), pandas (v1.3.5), parasail (v1.3.3), polars (v0.18.3), pytorch (v1.12.1), pytorch-metric-learning (v2.3.0), scikit-learn (v1.0.2), seaborn (v0.12.2), tqdm (v4.65.0). The following tools were used for data processing and analysis: guppy (v6.3.8), minimap2 (v2.25), modkit (v0.2.0). Nextflow pipeline for Illumina raw data alignment alignment can be found at https://github.com/UMCUGenetics/NF-IAP. The pipeline for variant filtering can be found at https://github.com/ToolsVanBox/SMuRF.

## Acknowledgements

We acknowledge the Utrecht Sequencing Facility (USEQ) for providing sequencing service and data. USEQ is subsidized by the University Medical Center Utrecht and The Netherlands X-omics Initiative (NWO project 184.034.019). We thank Edwin C. A. Stigter and Mehmet C. Gülersönmez for their help on mass-spectrometry validation of the synthetic oligos.

## Contributions

M.P.-G., T.B.D., B.M.T.B. and J.d.R. conceived the project. M.P.-G developed the 8-oxo-dG caller. M.P.-G and N.J.M.B performed the experiments to ligate the synthetic oligos. D.M.K.v.S performed the cell lines experiments. J.P.K and M.J.v.R performed analysis on Illumina derived mutational signatures. A.M. provided feedback on oligo design. R.S and C.V provided feedback in algorithm implementations. R.v.B provided feedback on mutational analysis. M.P.-G drafted the first version of the manuscript with guidance from J.d.R, T.B.D and B.M.T.B. J.d.R, T.B.D and B.M.T.B contributed to major parts of the manuscript and revised the manuscript. All authors read and approved of the final manuscript.

## Competing interests

J.d.R and A.M are co-founders and directors of Cyclomics, a genomics company, they declare no competing interests. M.P-G, D.M.K.v.S, N.J.M.B, R.S, J.P.K, C.V, M.J.v.R, R.v.B, B.M.T.B, and T.B.D declare no competing interests.

## Funding

This work was funded by Health Holland (No. LSHM19029). B.M.T.B, J.d.R and R.v.B are members of the Oncode Institute, which is partly funded by the Dutch Cancer Society (KWF Kankerbestrijding). T.B.D was funded by the Dutch Cancer Society (KWF Kankerbestrijding, KWF grant 14798).

## Supplementary material

### List of Supplementary Notes

**Supplementary note 1: 8-oxo-dG signal alignment.** The measured to expected signal alignment is not optimal because the expected signal of 8-oxo-dG is unknown. We use the expected signal of G instead, which will likely produce misassignment of data points to neighboring bases to optimize for the minimum distance between the two sequences. For example, as seen in **Figure 1c**, the signal produced by 8-oxo-dG is clearly different from G, but the number of measurements in the 8-oxo-dG and previous G bases are low compared to the rest.

**Supplementary note 2: Unsuccessful strategies.** We explored whether we could increase the specificity by filtering potential false positive calls. For example, the *Remora* model depends on accurate G calls from the Bonito model. We therefore evaluated if inaccurate G calls from Bonito would lead to FP 8-oxo-dG calls in the *Remora* model. However, we observed that filtering miscalls had a minimal impact (0.1%) in 8-oxo-dG FP reduction. We also explored the use of metric learning in an effort to facilitate the learning process of the model by defining triplets of examples in which the 5-mer context of the examples was the same. However, this strategy quickly led to an overfitting of the model, and was deemed non-viable (**Supplementary figure 39, Supplementary figure 40**)

**Supplementary note 3: Chromosomal variability of 8-oxo-dG levels.** We evaluated whether the observed differences in 8-oxo-dG levels across chromosome arms could be explained by their 5-mer content, and fitted a linear model with 5-mer relative abundances as independent variables and 8-oxo-dG levels as response variables. However, none of the regression coefficients was significant based on a permutation test (**Supplementary figure 41**), indicating that sequence alone does not explain these differences in 8-oxo-dG levels. Since we observed different 8-oxo-dG levels per genomic region, we then hypothesized whether these could explain the observed chromosomal variability. We evaluated if the relative abundance of the different genomic regions could explain the observed chromosomal differences. However, we obtained a similar result, with none of the coefficients being significant (**Supplementary figure 42**).

**Supplementary note 4: Telomere ligation.** DNA molecules prepped for nanopore sequencing require the ligation of sequencing adapters that contain the motor protein that will thread the DNA molecule through the pore. Adapter ligation can only be successful on a blunt end of a double strand DNA molecule. Telomere DNA has a long overhang to prevent chromosome unwinding. This overhang prevents the adapter ligation, unless the opposite strand is elongated to the length of the overhang [55, 67]. That means that the forward strand (p-arm) and reverse strand (q-arm) cannot be sequenced using nanopore sequencing, unless a break occurs somewhere within the double stranded sequence of the telomere. On the other hand, the reverse strand (p-arm) and forward strand (q-arm), can be sequenced when a break occurs somewhere within the chromosome as long as the DNA molecule is long enough. The latter is more likely given the relative short length of telomeres compared to the rest of the chromosome.

### List of Supplementary Figures

**Supplementary figure 1:**
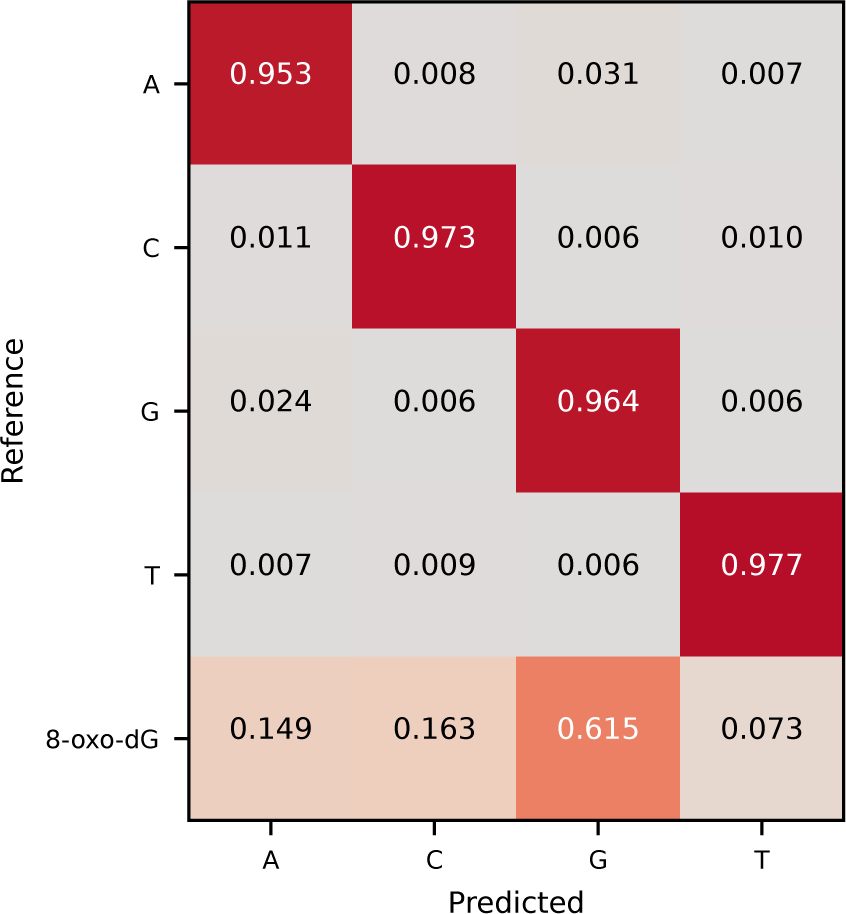
8-oxo-dG error rate ONT pre-trained *Bonito*. Confusion matrix of the ONT Bonito pre-trained model evaluated on the test fold of the 8-oxo-dG containing oligo dataset. Values indicate the fraction of outcomes for each ground truth base.

**Supplementary figure 2:**
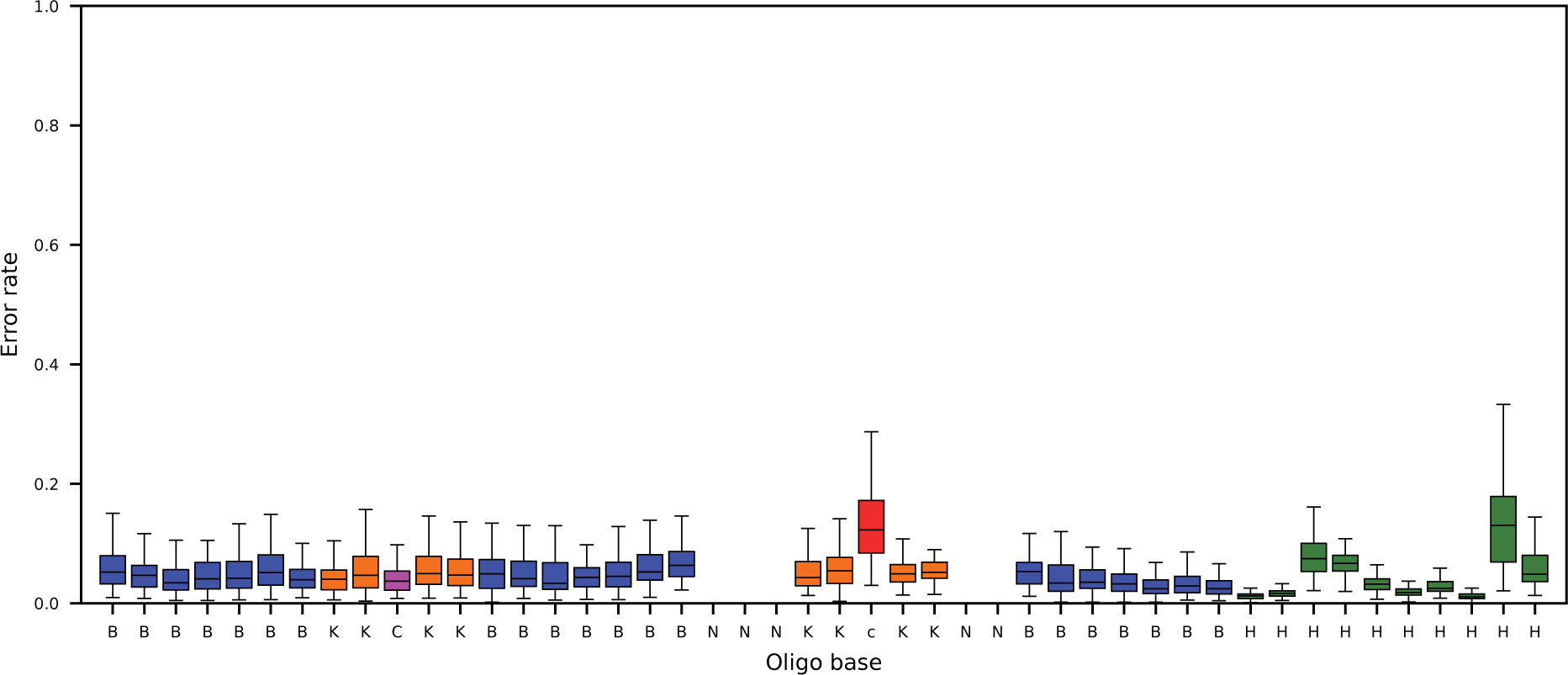
Reverse oligo strand error rate. Error rate per oligo base across all sequenced non-8-oxo-dG containing repeats. Random bases are excluded from the analysis as we do not know their true reference.

**Supplementary figure 3:**
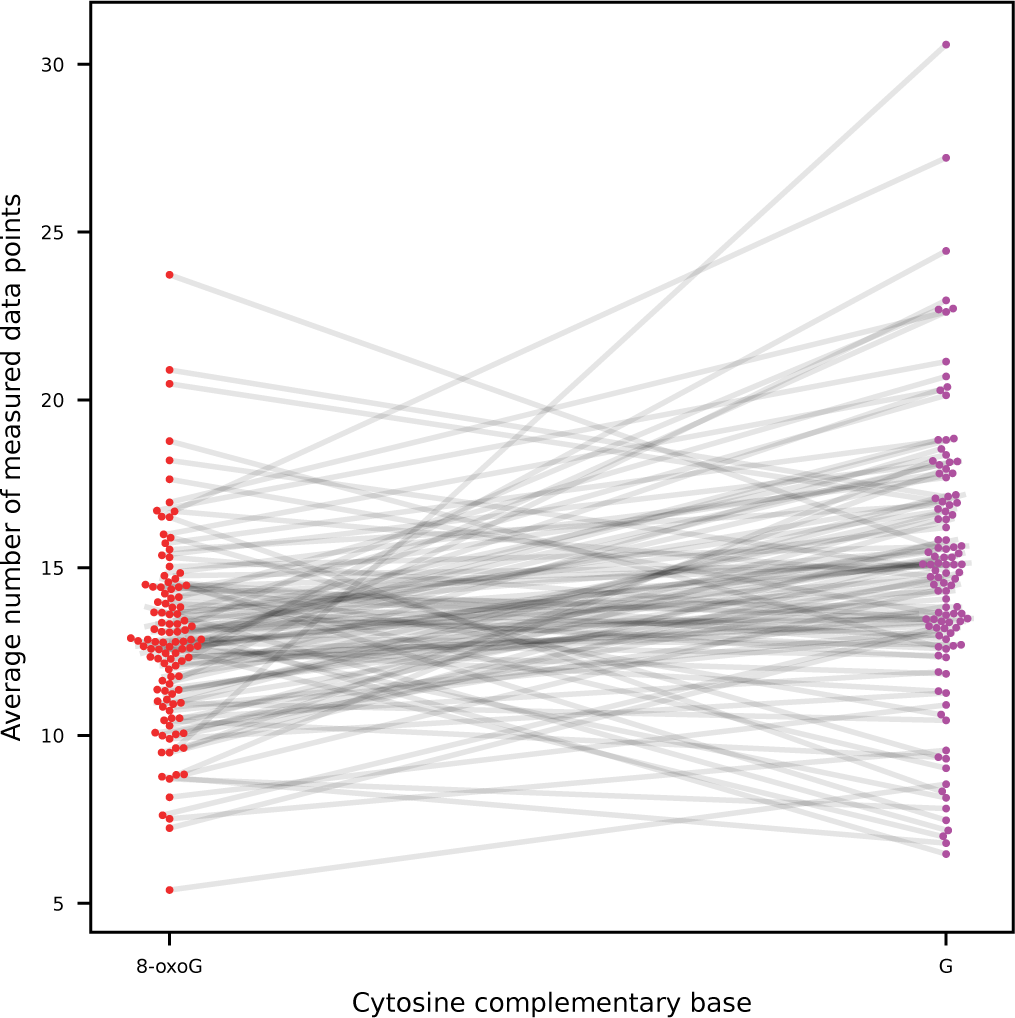
Cytosine speed changes, 8-oxo-dG vs G. Average number of measured data points (as segmented using Tombo) per cytosine (n=1000) when paired with 8-oxo-dG (red) or G (purple). Cytosines in the same 5-mer context (KKCKK) are connected via gray lines.

**Supplementary figure 4:**
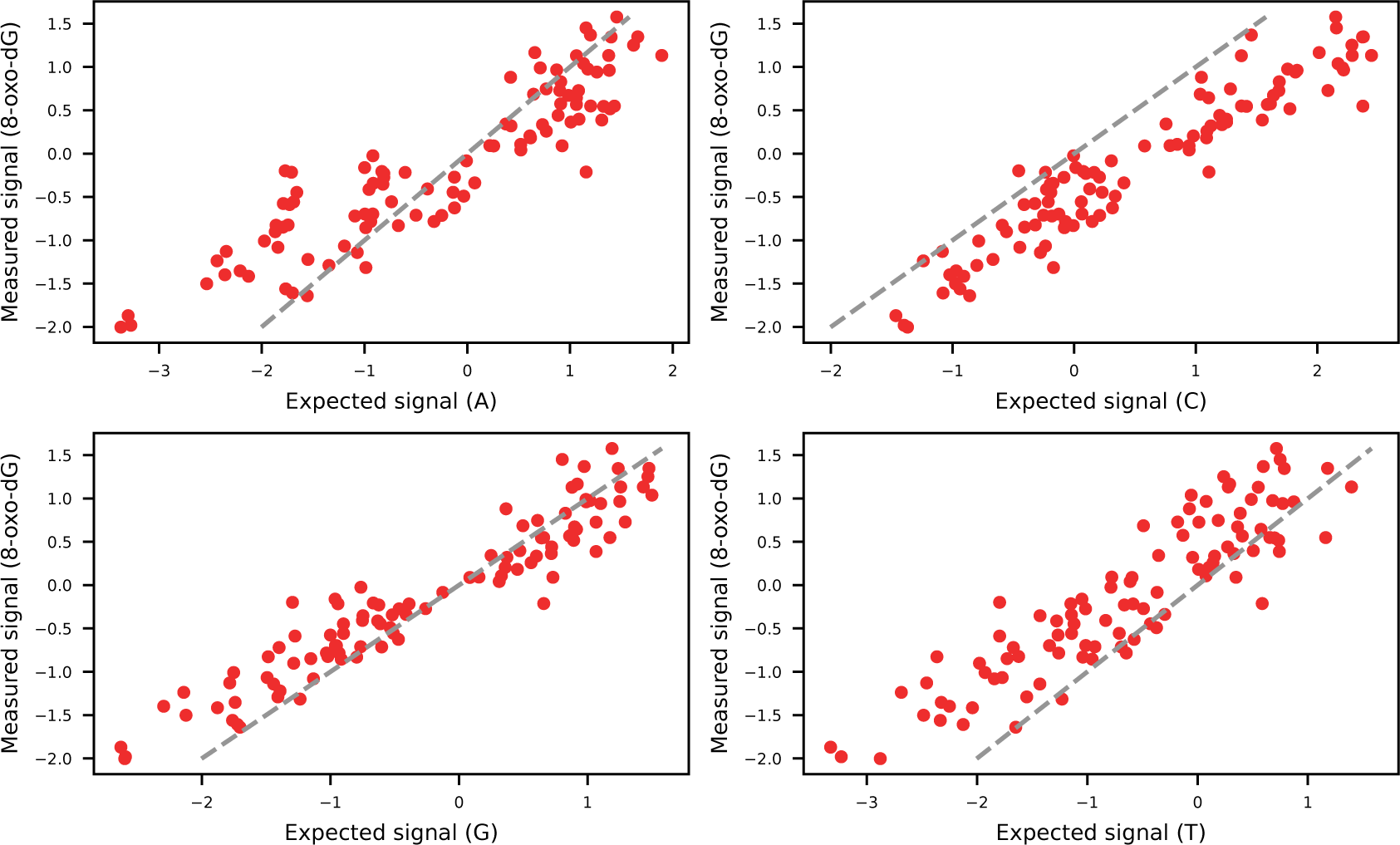
**8-oxo-dG versus canonical bases signals.**Average measured normalized canonical base (A, C, G, T) signal and 8-oxo-dG signal per measured 5-mer as segmented using Tombo. Identity line indicated as the dashed gray line.

**Supplementary figure 5:**
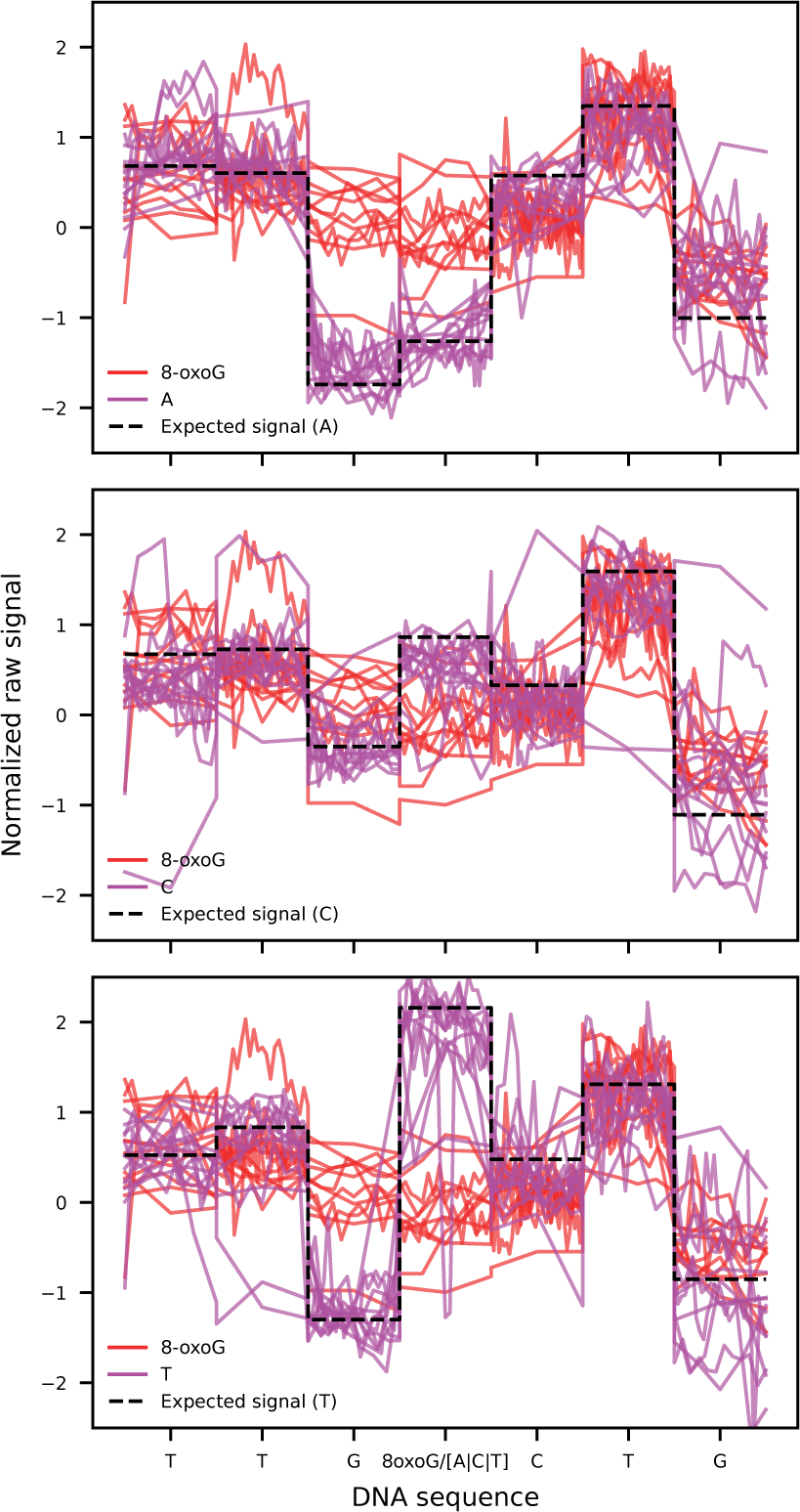
8-oxo-dG signal examples. Example of 8-oxo-dG (red) and A|C|T (purple) signals in the TTG(8-oxo-dG/A|C|T)CTG context. Dashed black line indicates the expected signal value based on the A|C|T containing sequence.

**Supplementary figure 6:**
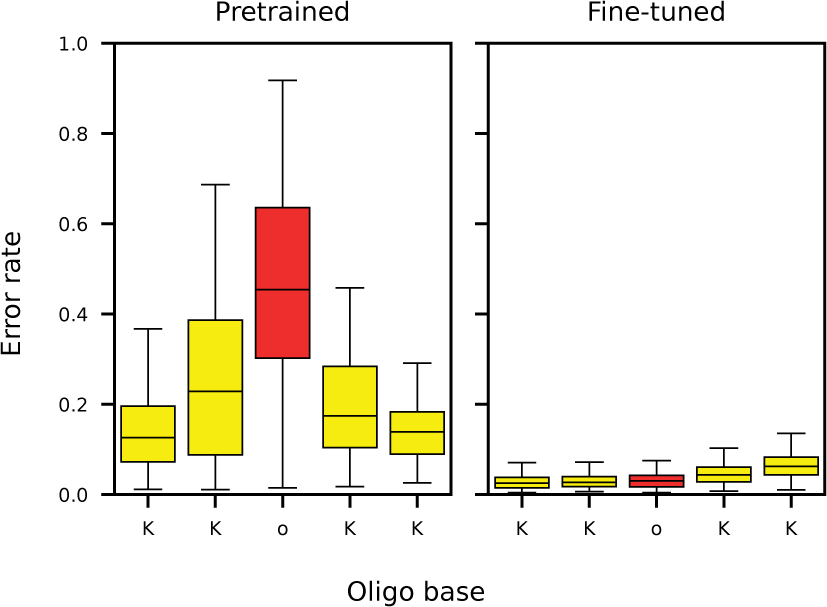
**8-oxo-dG error rate after *Bonito* fine-tuning.**Error rate per base around 8-oxo-dG (o) in the ONT *Bonito* pre-trained model (left), and fine-tuned (right) model with both T2T and oligo data. Error rate includes mismatches and deletions.

**Supplementary figure 7:**
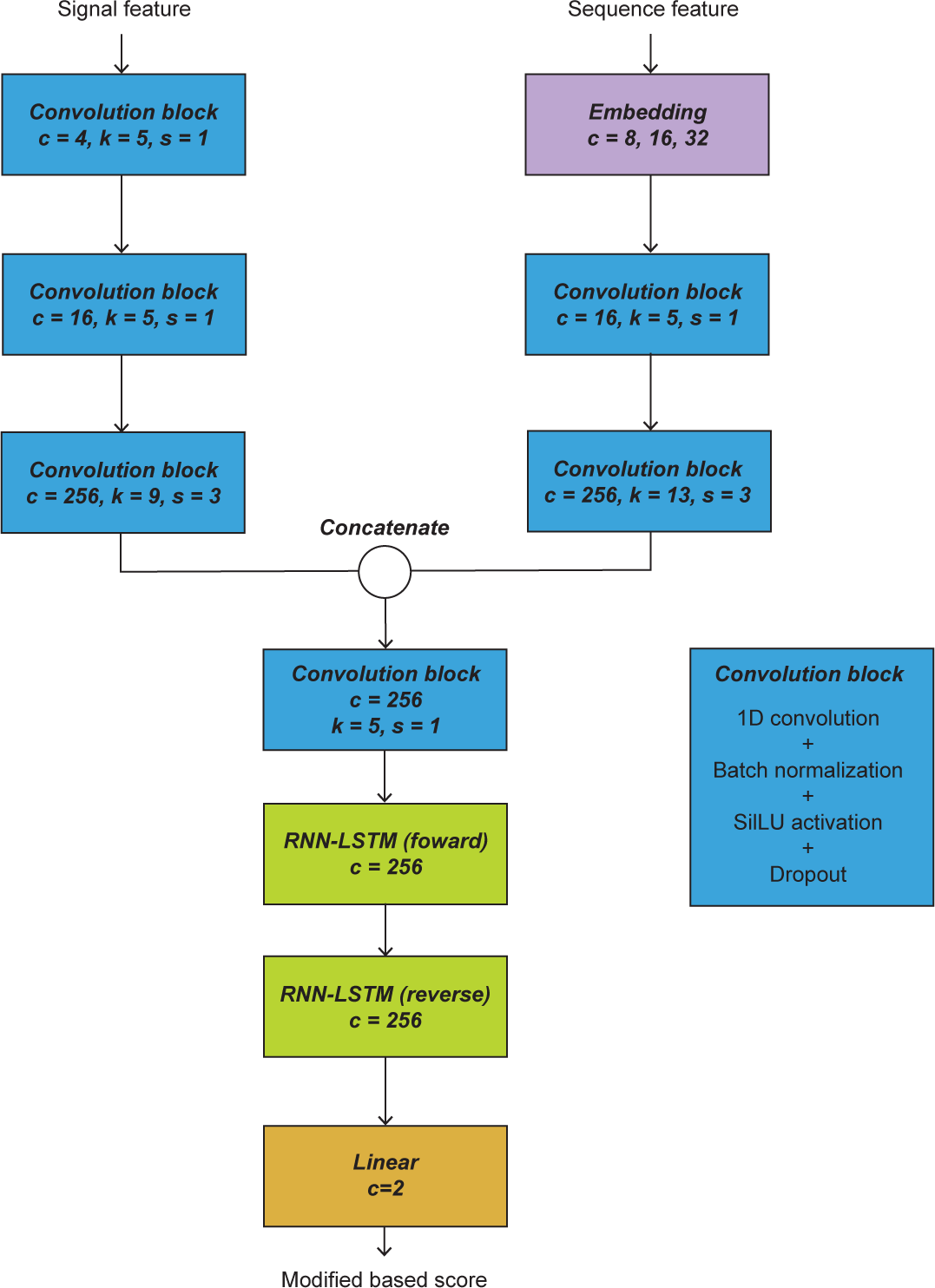
*Remora* model architecture. Schematic representation of the neural network architecture for a *Remora* model. Numbers indicate output dimension (c), kernel size (k), stride (s).

**Supplementary figure 8:**
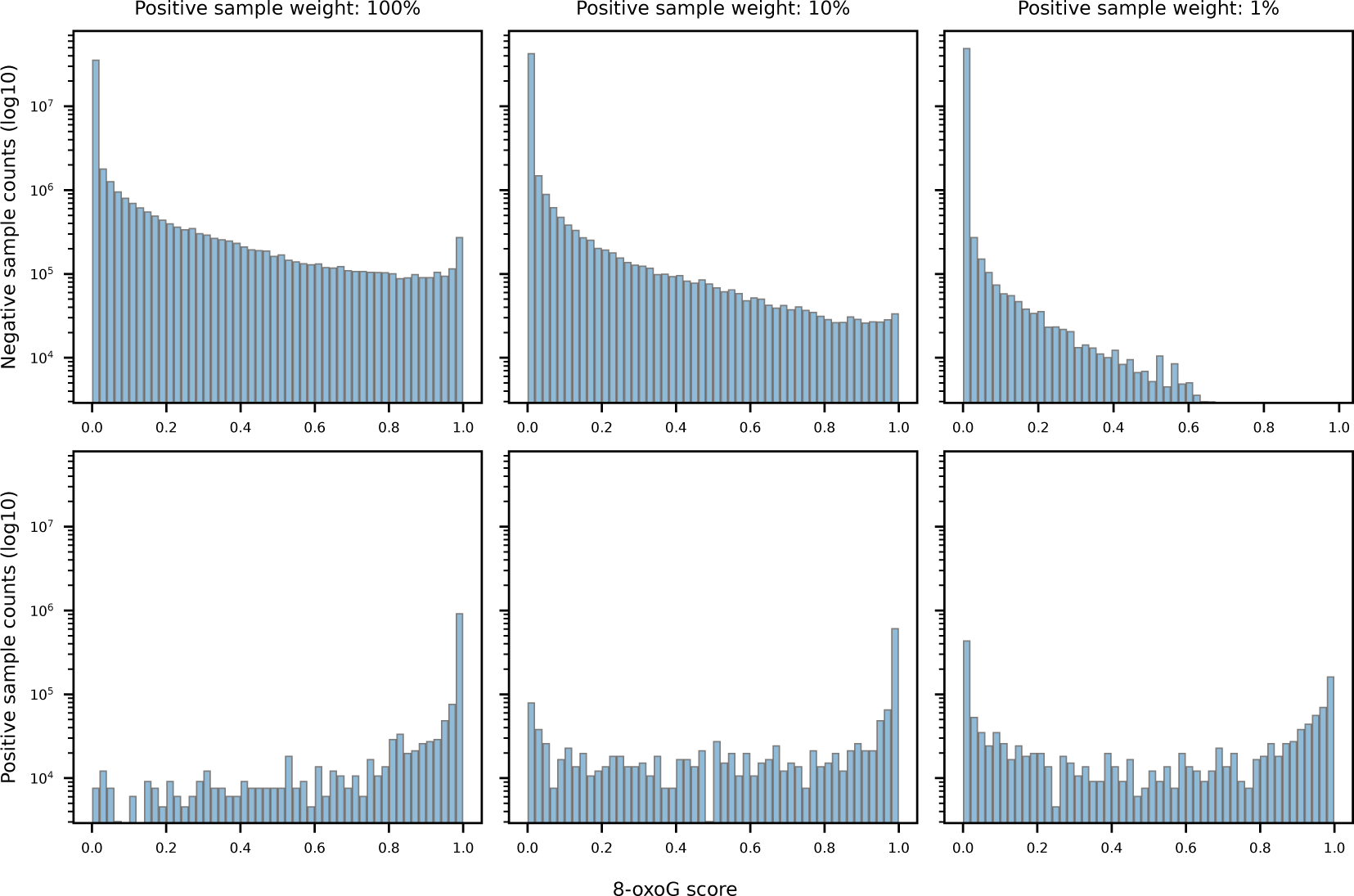
*Remora* model classification score distributions. Score distributions for the negative (top) and positive (samples) of the test fold, for each trained *Remora* base model with different positive sample weights: 100%, 10% and 1% from left to right.

**Supplementary figure 9:**
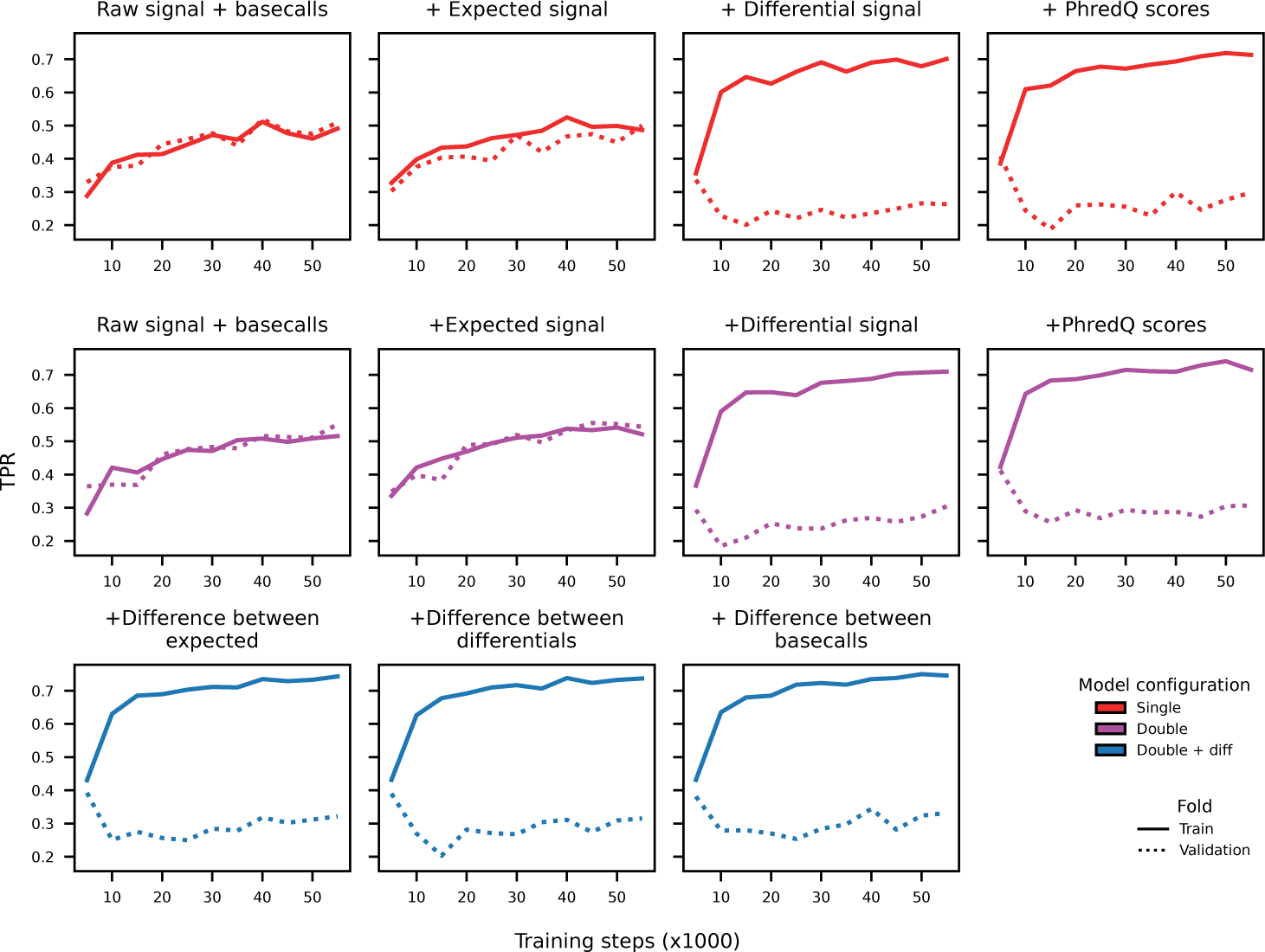
*Remora* model during training performance with increased number of features. True positive rate overt training of a *Remora* model with different features. Each model contains one additional feature from top left to bottom right.

**Supplementary figure 10:**
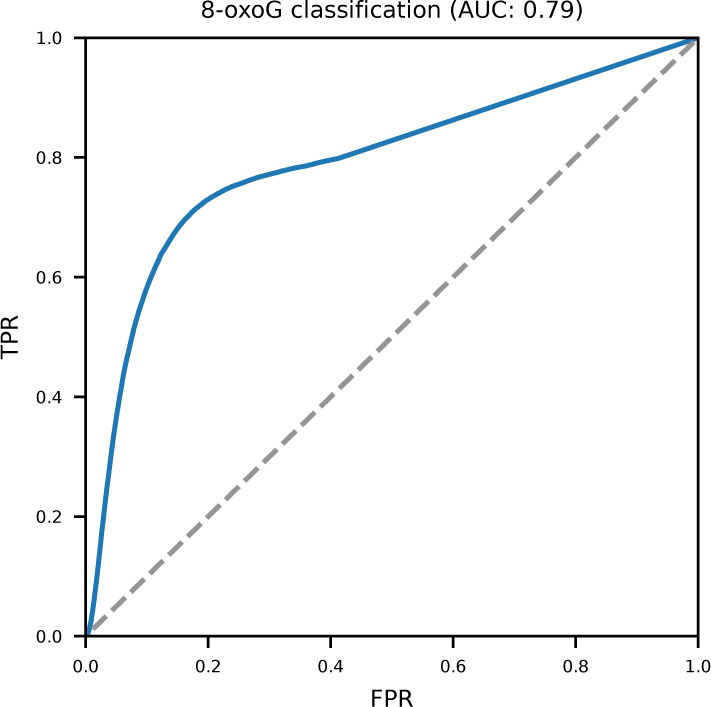
Single feature classifier. ROC curve (blue line) for a classifier solely based on the absolute sum of differences between measured raw signal and expected signal (Finetuned - Pretrained). This feature is highly differentiating since it achieves by itself an AUC of 0.79. Dashed gray line indicates the performance of a random classifier.

**Supplementary figure 11:**
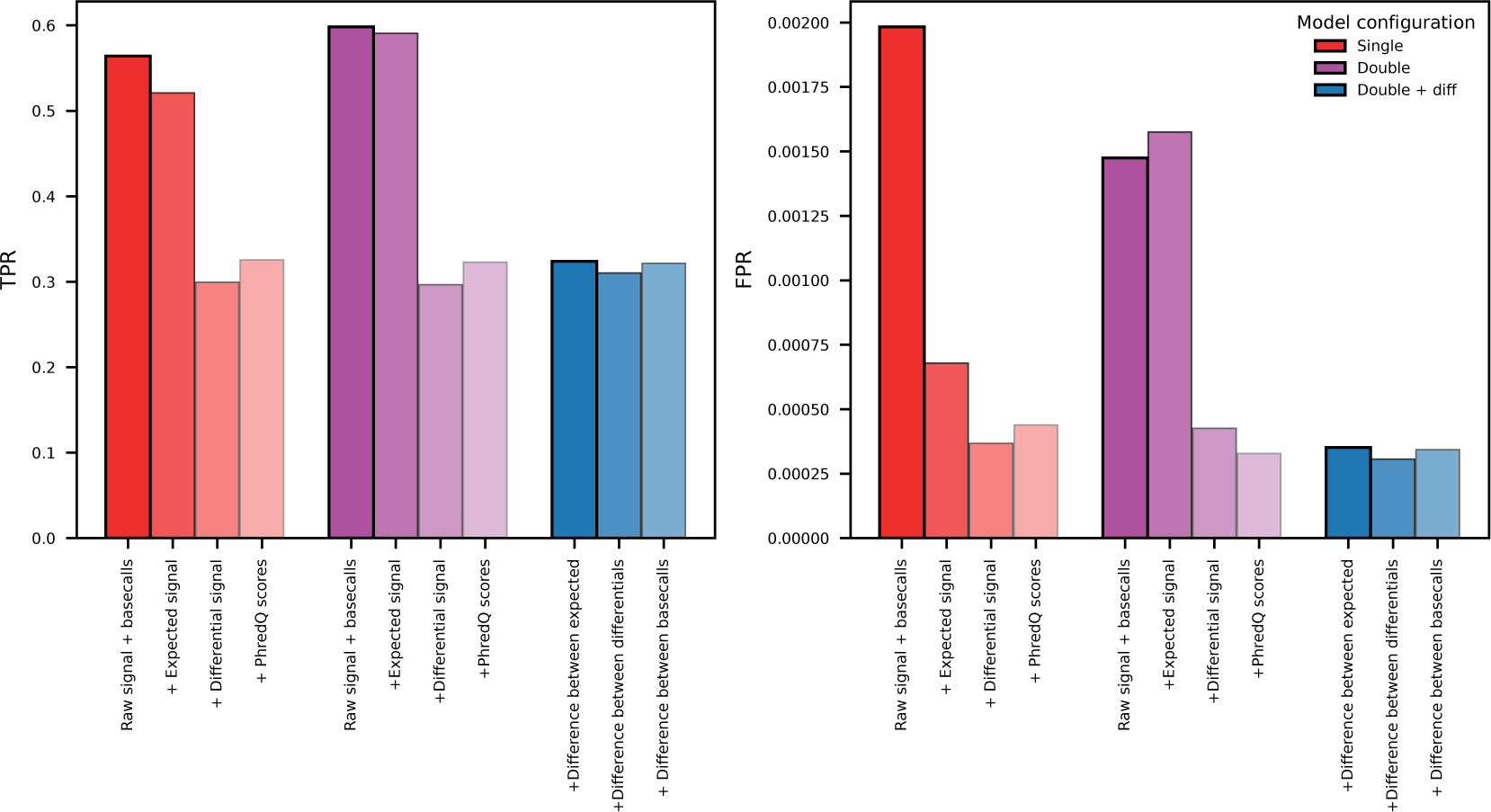
*Remora* model performance with additional features. TPR and Q-Score specificity evaluated on the test fold for the experiment in which additional features were added sequentially. Metrics are calculated using a 0.5 threshold. Models include additional features in a cumulative manner, from left to right: basecalls, expected signal, difference between expected and measured signal, and basecall phred quality scores. Red bars include features from the fine-tuned model, purple bars also include features from the *Bonito* pre-trained model, blue bars include the difference between the features of the *Bonito* fine-tuned and pre-trained models.

**Supplementary figure 12:**
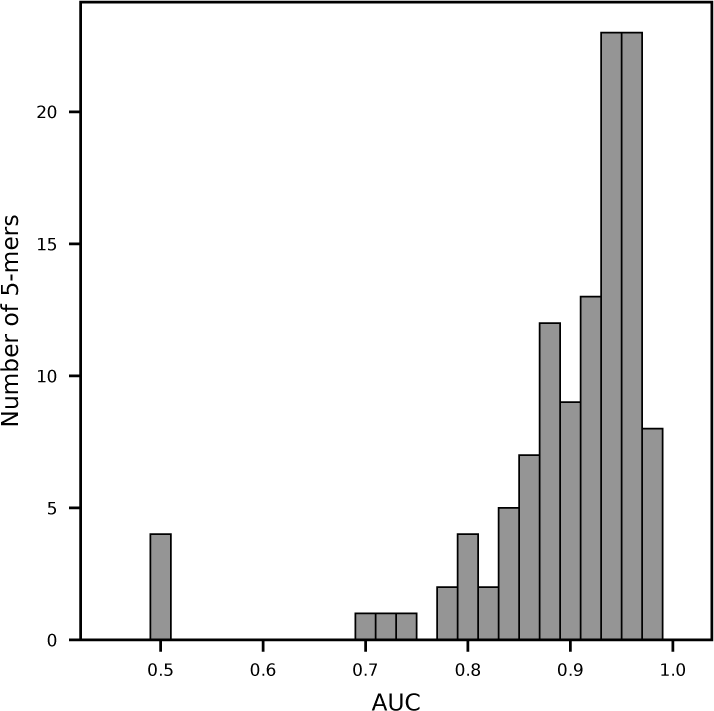
AUC per 5-mer. 8-oxo-dG classification performance for each 5-mer as area under the curve (AUC).

**Supplementary figure 13:**
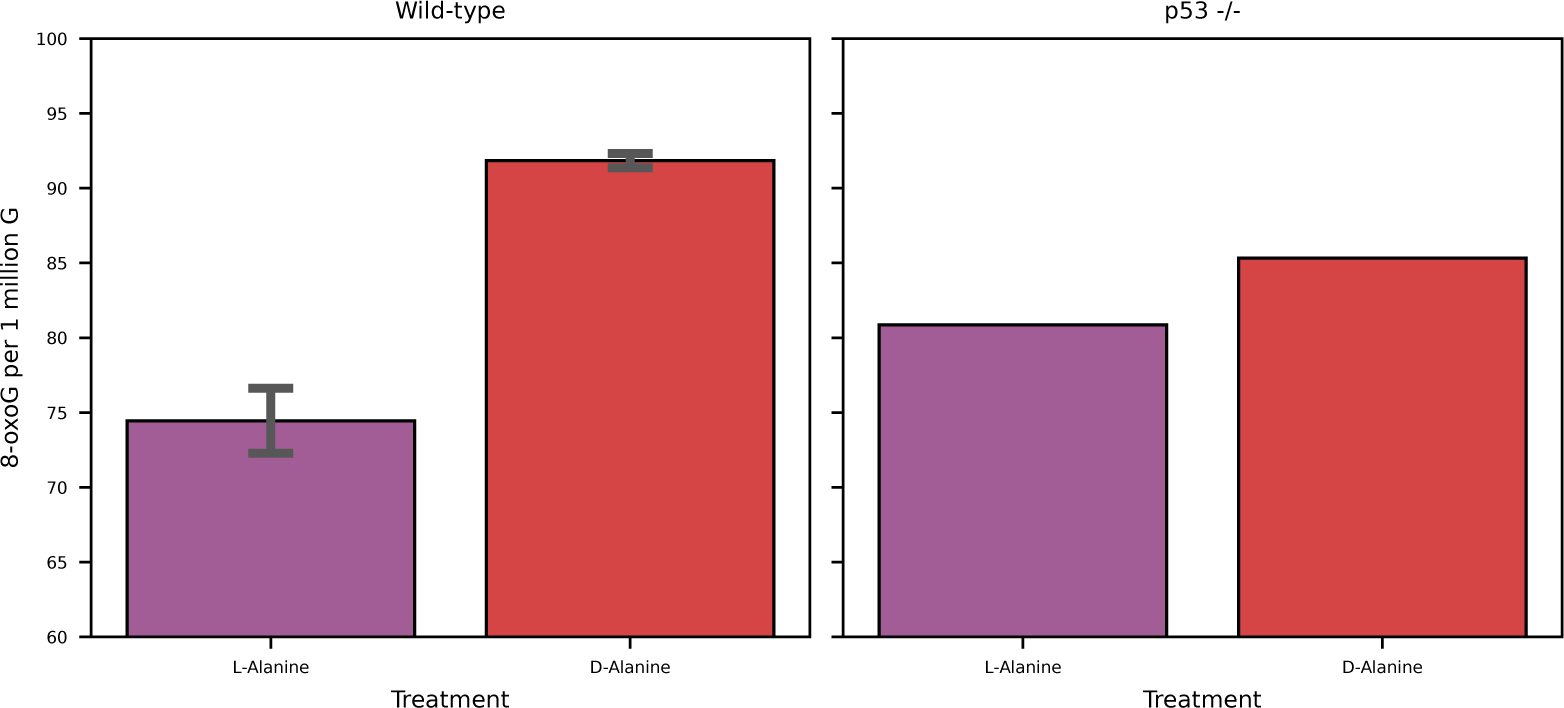
8-oxo-dG levels per cell line and treatment. Overall 8-oxo-dG molecules per 1 million G molecules per L-Alanine and D-Alanine treated cells. Error bars indicate minimum and maximum calculated values per biological replicate.

**Supplementary figure 14:**
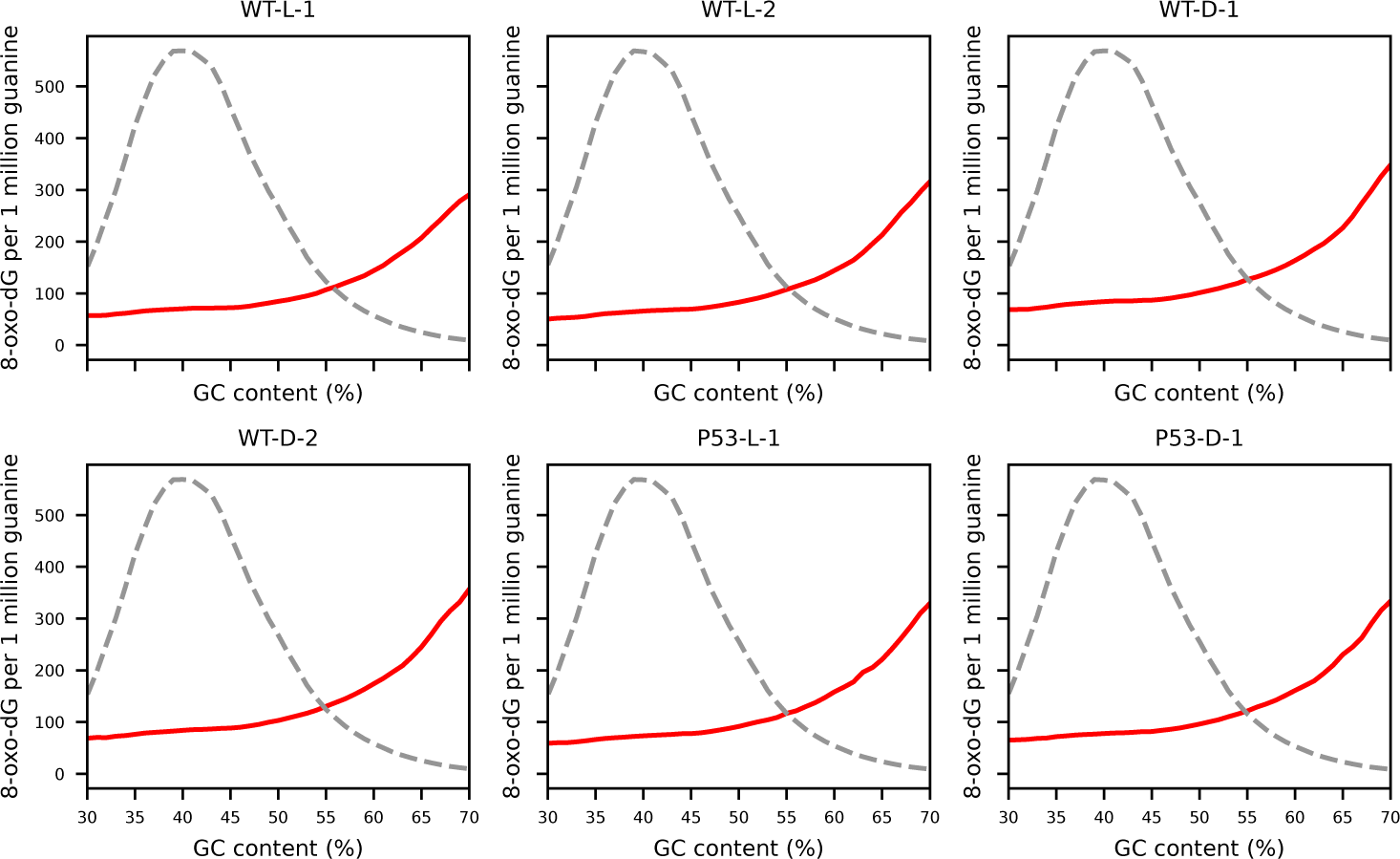
**8-oxo-dG levels per GC content.**8-oxo-dG levels across different GC (%) content bins. Red lines indicate 8-oxo-dG values per 1 million guanines. Grey dashed line indicates the distribution of measured GC content bins.

**Supplementary figure 15:**
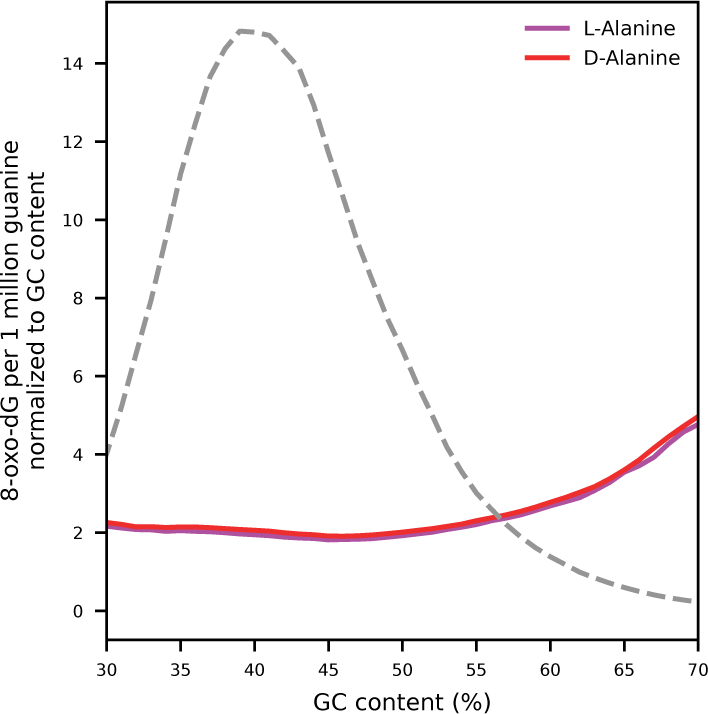
8-oxo-dG levels normalized to GC content. 8-oxo-dG levels across different GC (%) content bins. Purple (L-Alanine) and red (D-Alanine) lines indicate 8-oxo-dG values per 1 million guanines normalized to GC content. Grey dashed line indicates the distribution of measured GC content bins.

**Supplementary figure 16:**
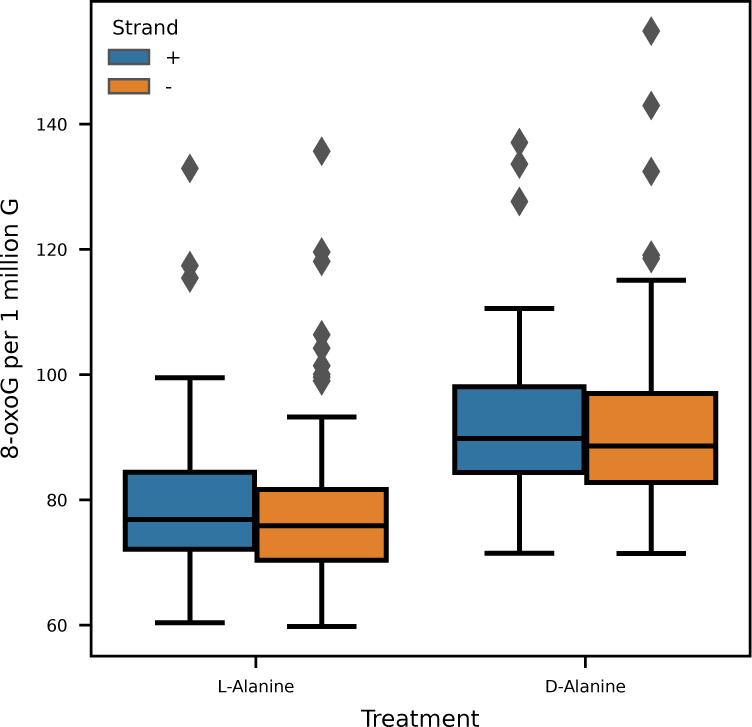
8-oxo-dG levels normalized to GC content. 8-oxo-dG counts per 1 million guanine divided per DNA strand and L-alanine or D-alanine treated cells. Boxplot represents values per chromosome for all cell lines. Horizontal bar represents the median, boxes indicate the 25th and 75th percentiles, whiskers indicate the 10th and 90th percentiles, and diamonds indicate outliers.

**Supplementary figure 17:**
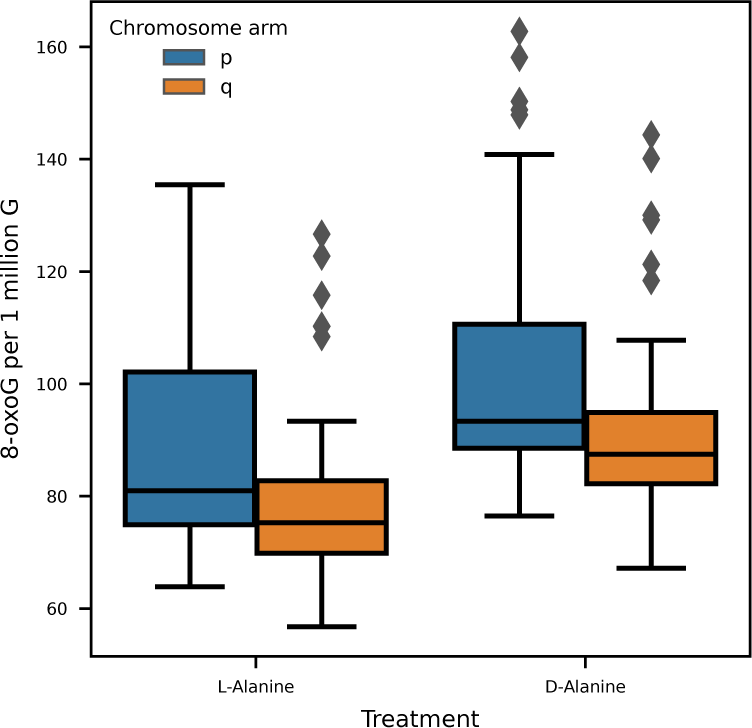
8-oxo-dG levels normalized to GC content. 8-oxo-dG counts per 1 million guanine divided per DNA chromosome arm and L-alanine or D-alanine treated cells. Boxplot represents values per chromosome for all cell lines. Horizontal bar represents the median, boxes indicate the 25th and 75th percentiles, whiskers indicate the 10th and 90th percentiles, and diamonds indicate outliers.

**Supplementary figure 18:**
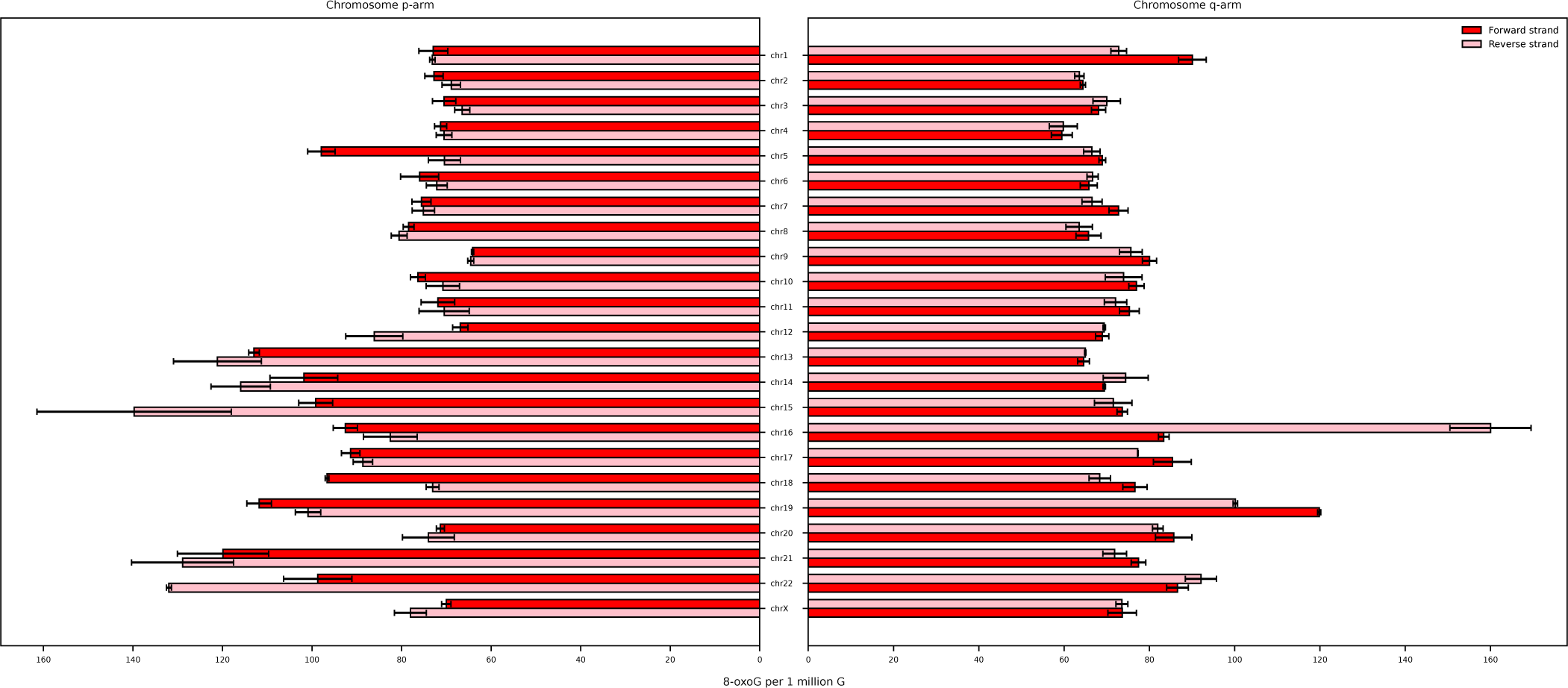
8-oxo-dG levels wild-type L-alanine treated cells. 8-oxo-dG counts per 1 million guanines for the RPE wild-type cells treated with L-alanine. Counts are divided per chromosome, chromosome arm and DNA strand. Error bars indicate the values of the two biological replicates.

**Supplementary figure 19:**
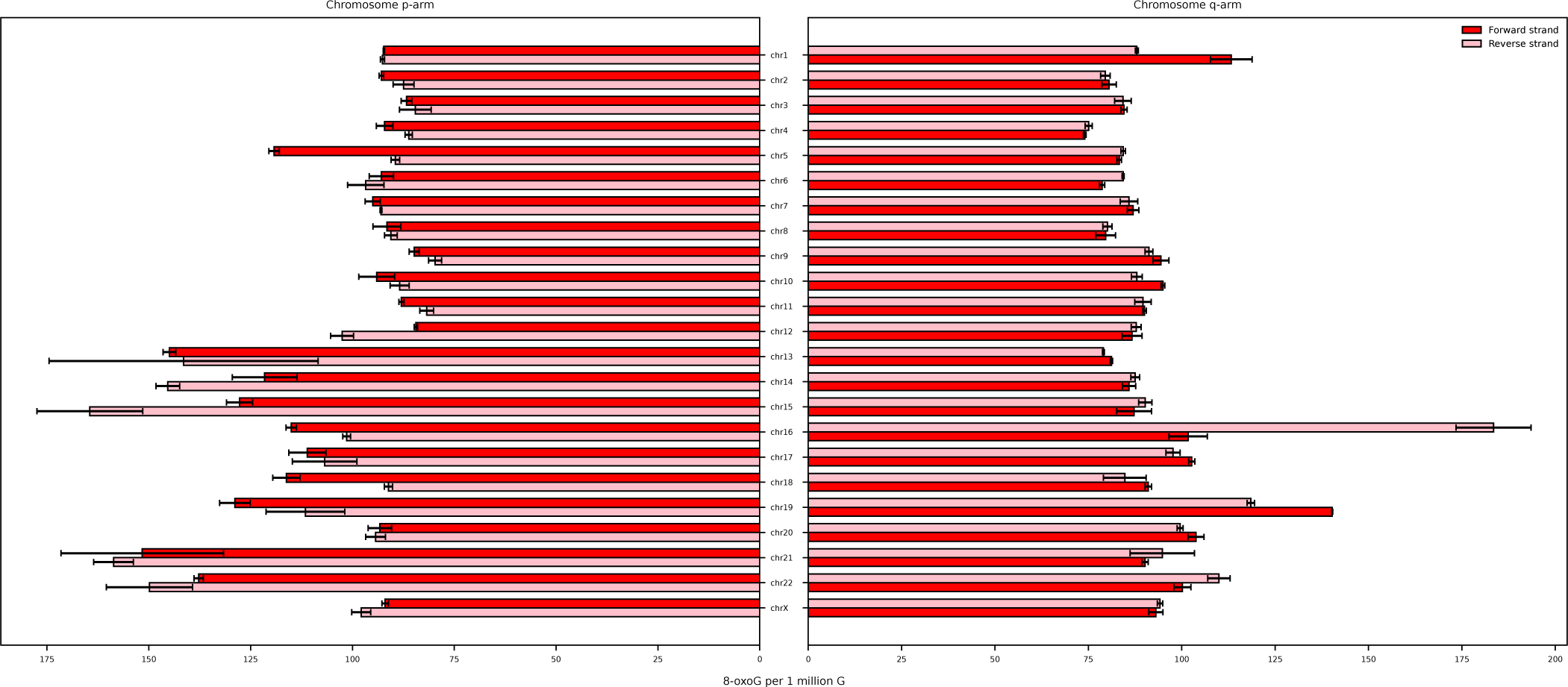
8-oxo-dG levels wild-type D-alanine treated cells. 8-oxo-dG counts per 1 million guanines for the RPE wild-type cells treated with D-alanine. Counts are divided per chromosome, chromosome arm and DNA strand. Error bars indicate the values of the two biological replicates.

**Supplementary figure 20:**
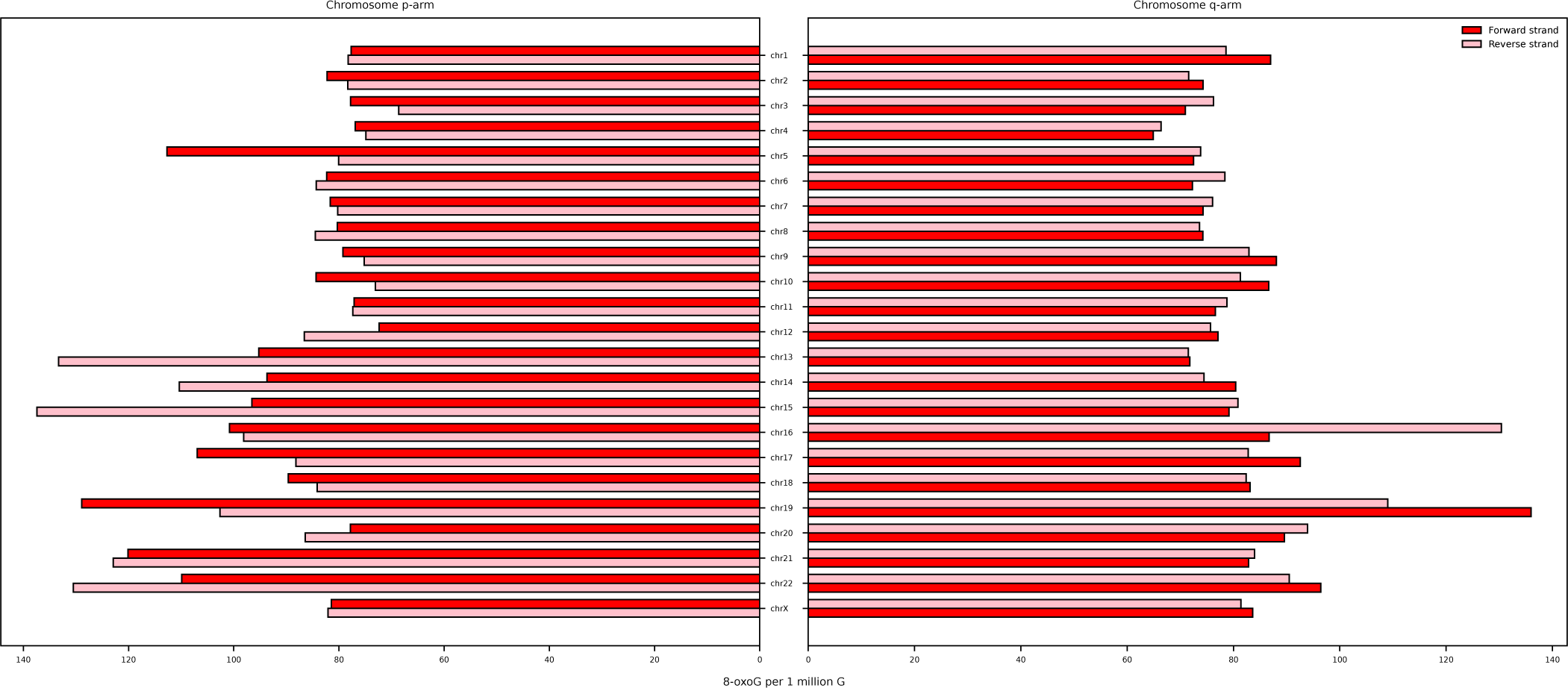
8-oxo-dG levels p53 KO L-alanine treated cells. 8-oxo-dG counts per 1 million guanines for the RPE wild-type cells treated with L-alanine. Counts are divided per chromosome, chromosome arm and DNA strand.

**Supplementary figure 21:**
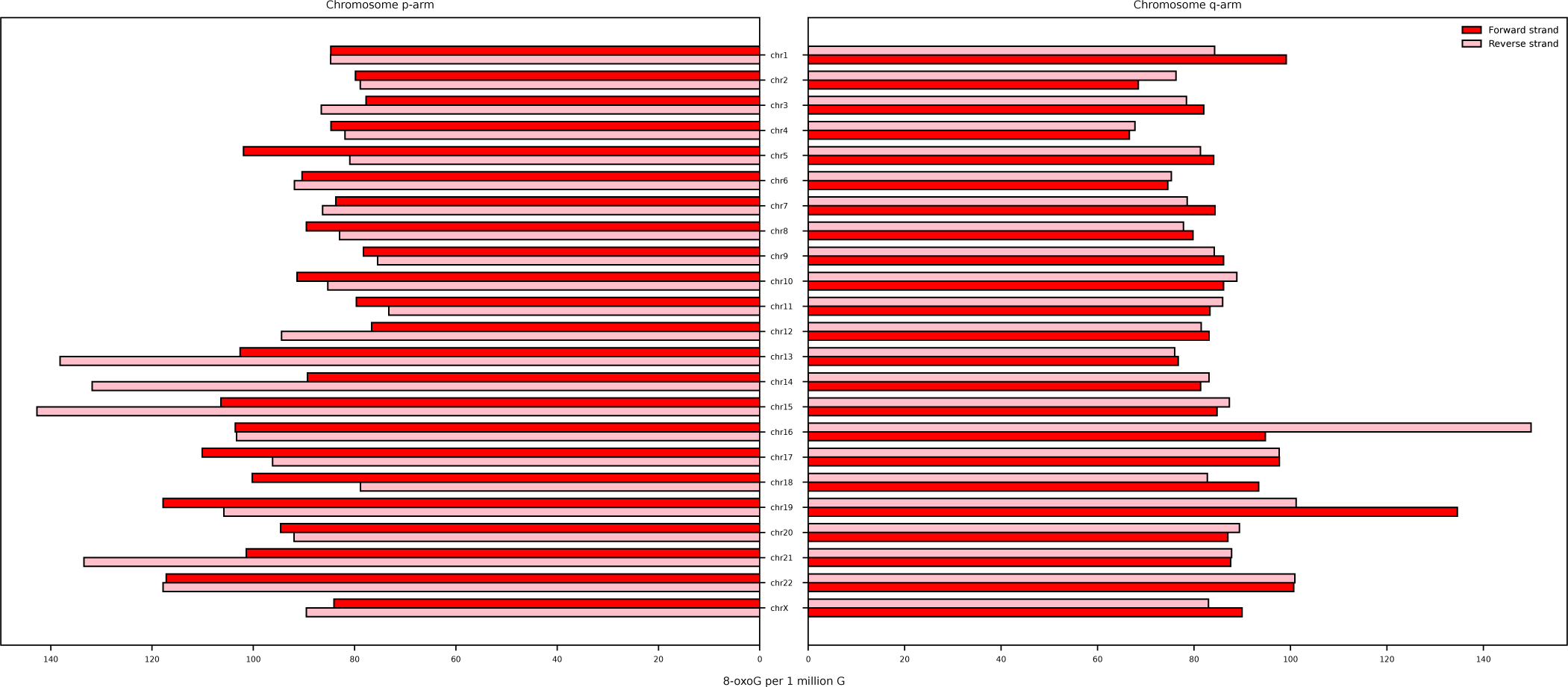
8-oxo-dG levels p53 KO D-alanine treated cells. 8-oxo-dG counts per 1 million guanines for the RPE wild-type cells treated with D-alanine. Counts are divided per chromosome, chromosome arm and DNA strand.

**Supplementary figure 22:**
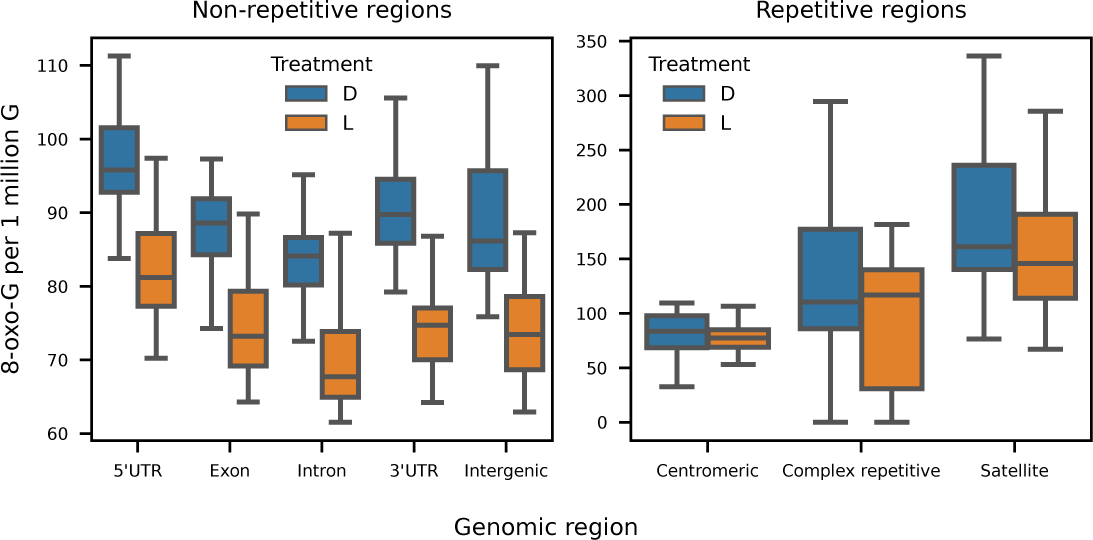
8-oxo-dG levels per genomic region and treatment. 8-oxo-dG counts per 1 million guanine grouped per genomic region based on the T2T reference genome assembly. Data values are collected per chromosome. Colors indicate whether cells were treated with D-Alanine (blue) or L-Alanine (orange). Horizontal bar represents the median, boxes indicate the 25th and 75th percentiles, whiskers indicate the 10th and 90th percentiles.

**Supplementary figure 23:**
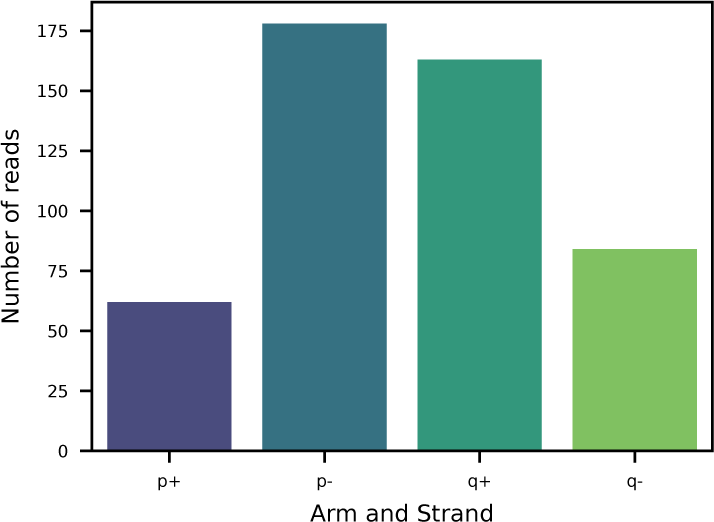
Number of reads mapped to telomeres. Number of mapped reads across all sequenced cell lines, split by chromosome arm and DNA strand. Only primarily mapped reads are included.

**Supplementary figure 24:**
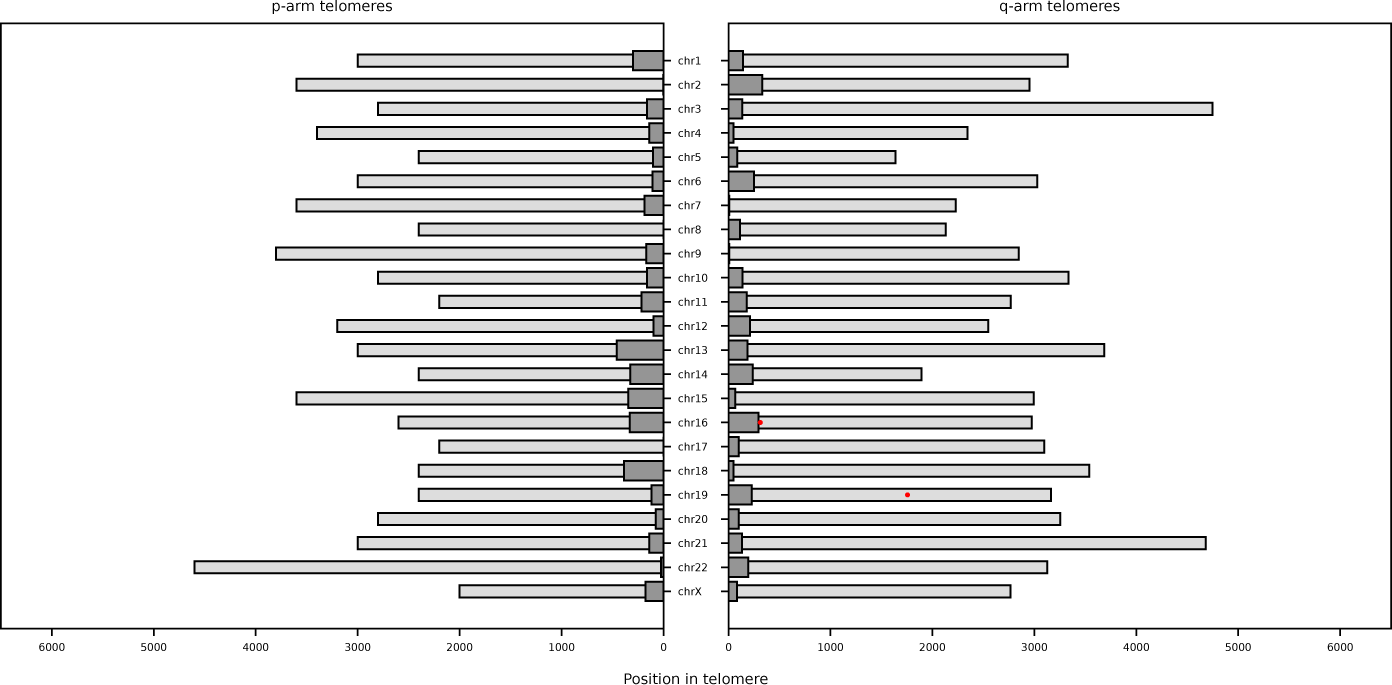
**8-oxo-dG molecules detected at telomeres.**8-oxo-dG calls (red) on reads that are primarily mapped to telomeric regions. Calls are combined from all the sequenced cell lines. Light gray bars indicate repetitive regions on each chromosome. Dark gray bars indicate non-repetitive regions, still annotated as telomeric regions on the T2T reference genome assembly.

**Supplementary figure 25:**
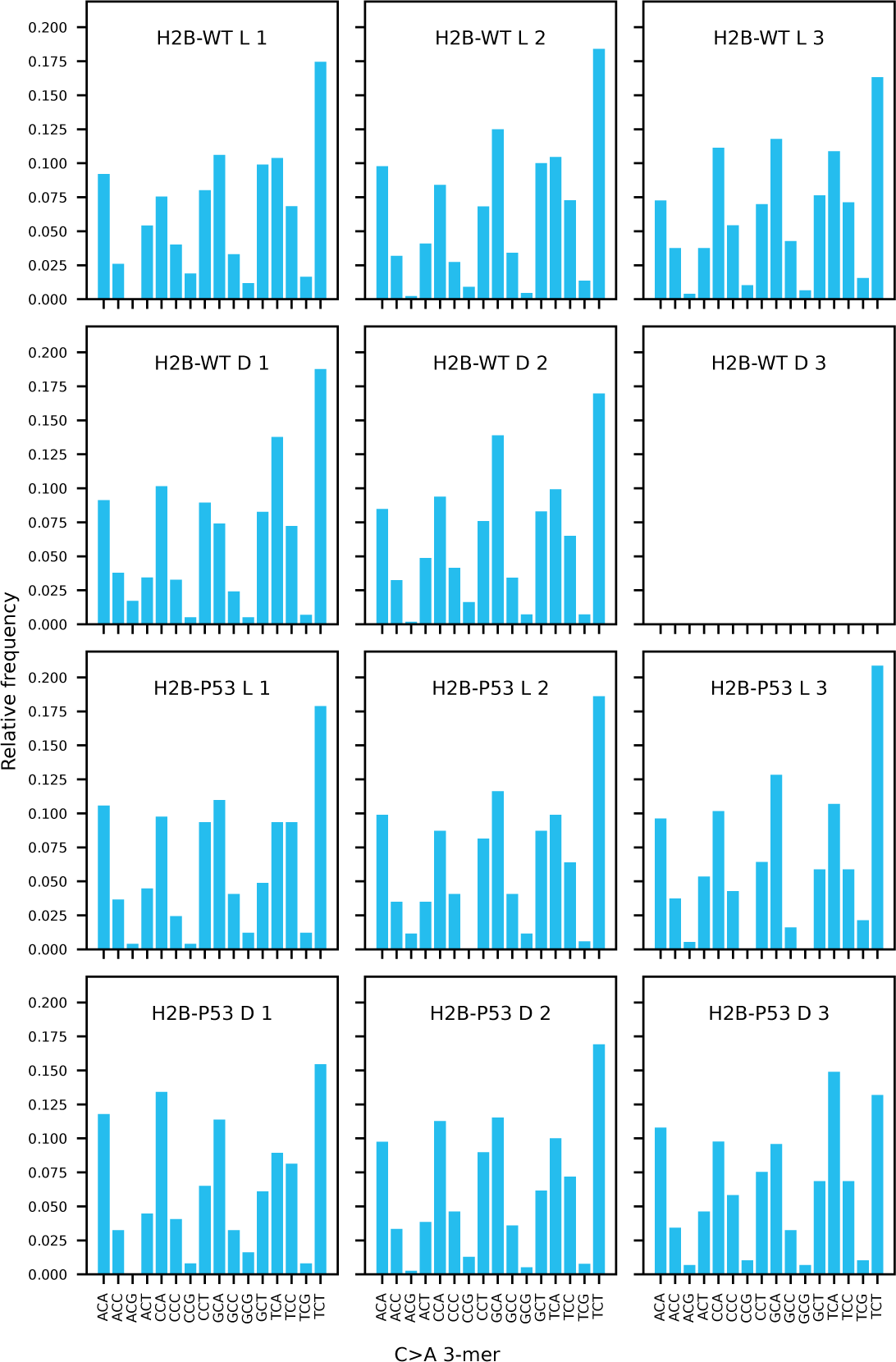
8-oxo-dG molecules detected at telomeres. Mutational profile for C>A mutations. Mutations are grouped based on their 3-mer context and counts are relative to the total amount of C>A mutations. Text in the plot indicates the cell line (WT or p53^-/-^), treatment (L-Alanine or D-Alanine), and the biological replicate number. H2B-WT-D3 was a failed sequencing run and therefore has no data.

**Supplementary figure 26:**
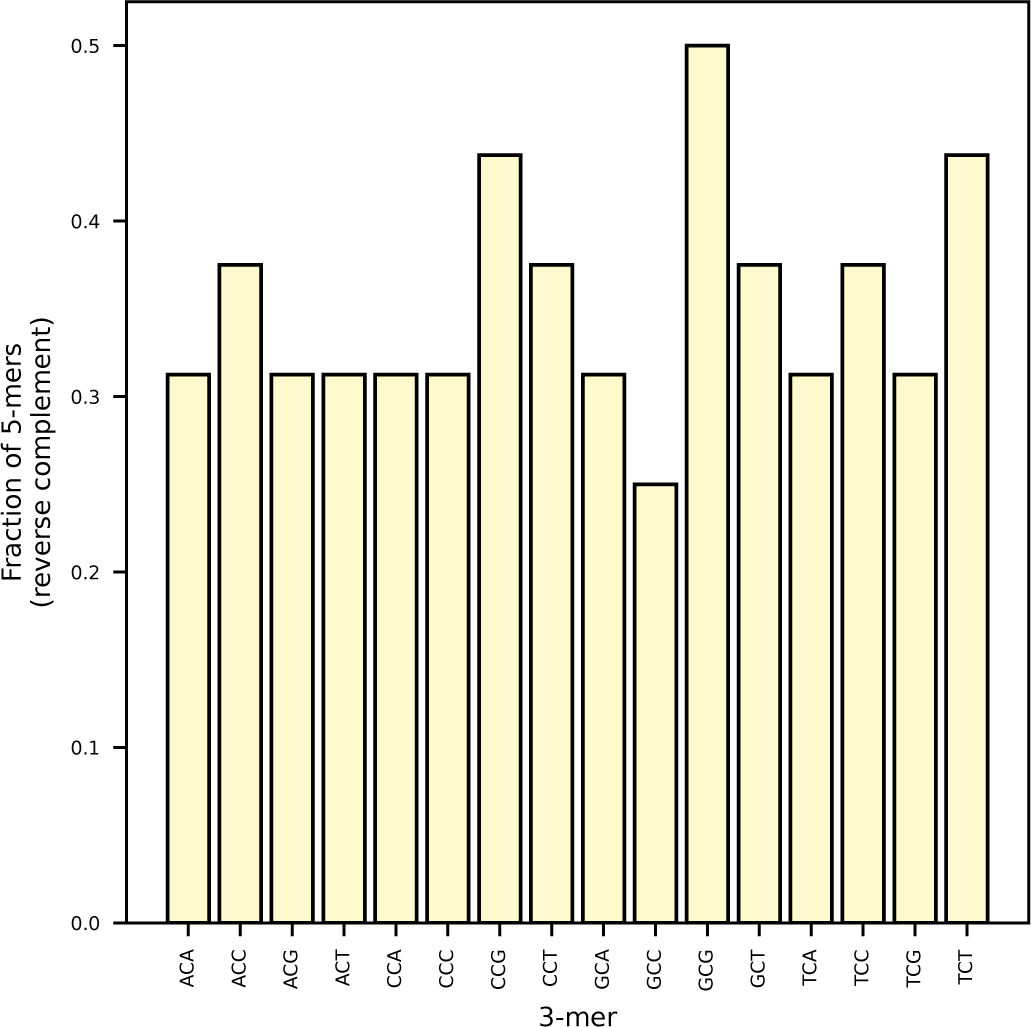
3-mer completness. Fraction of 5-mers with a specificity >Q40 for each 3-mer in the reverse complement strand.

**Supplementary figure 27:**
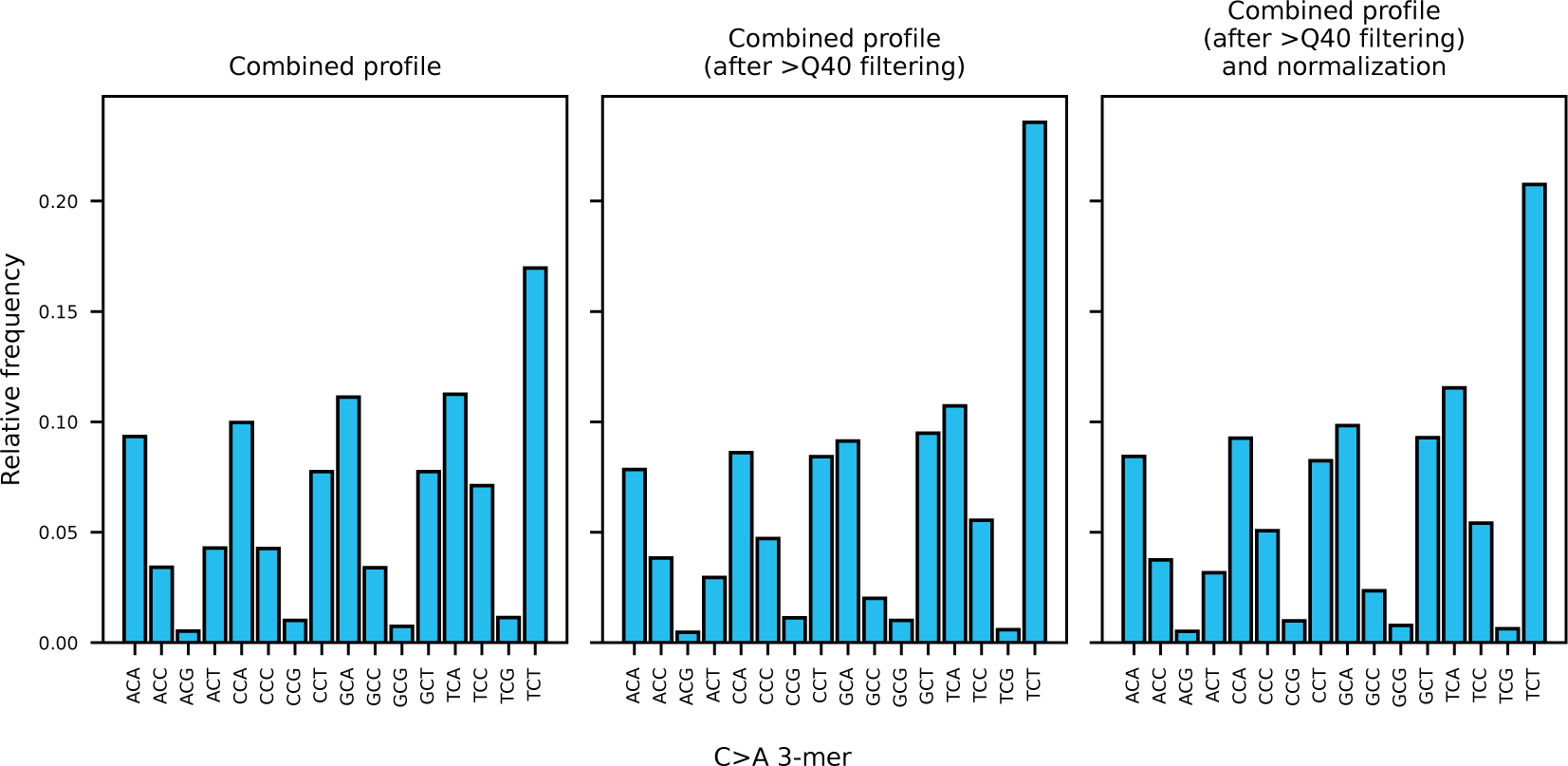
Mutational profile normalization. Mutational profile of C>A mutations of all cell lines and treatments joined. Left panel indicates the joined profiles without any correction. Middle panel includes only mutations in a 5-mer context with high specificity for 8-oxo-dG calling. Right panel includes corrections for the number of available 5-mers per 3-mer.

**Supplementary figure 28:**
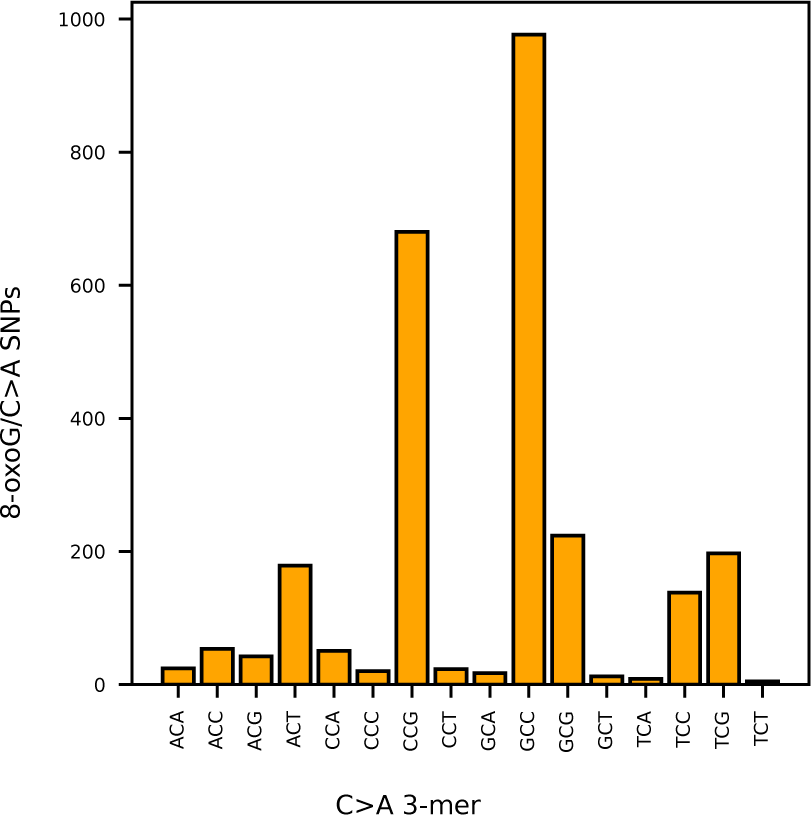
8-oxo-dG to mutation ratio. Ratio between 8-oxo-dG calls and C>A/G>T mutations in each C>A 3-mer. Includes data aggregated from all the cell lines.

**Supplementary figure 29:**
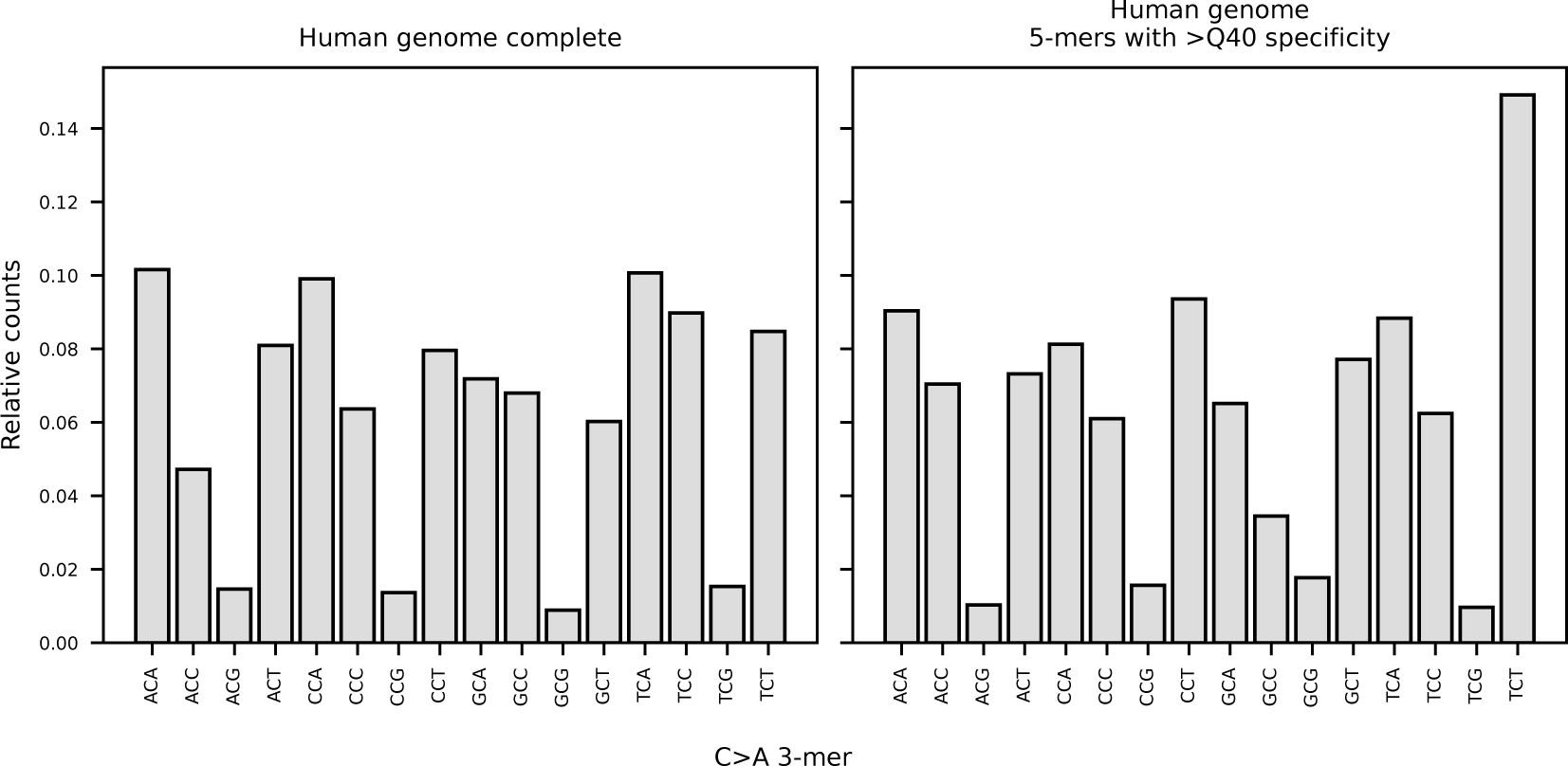
Human genome 3-mer content. Relative counts of all the 3-mers (with cytosine in the middle position) in the human genome. Left includes all the 3-mers, and right only includes counts of 5-mers with high specificity for 8-oxo-dG calling.

**Supplementary figure 30:**
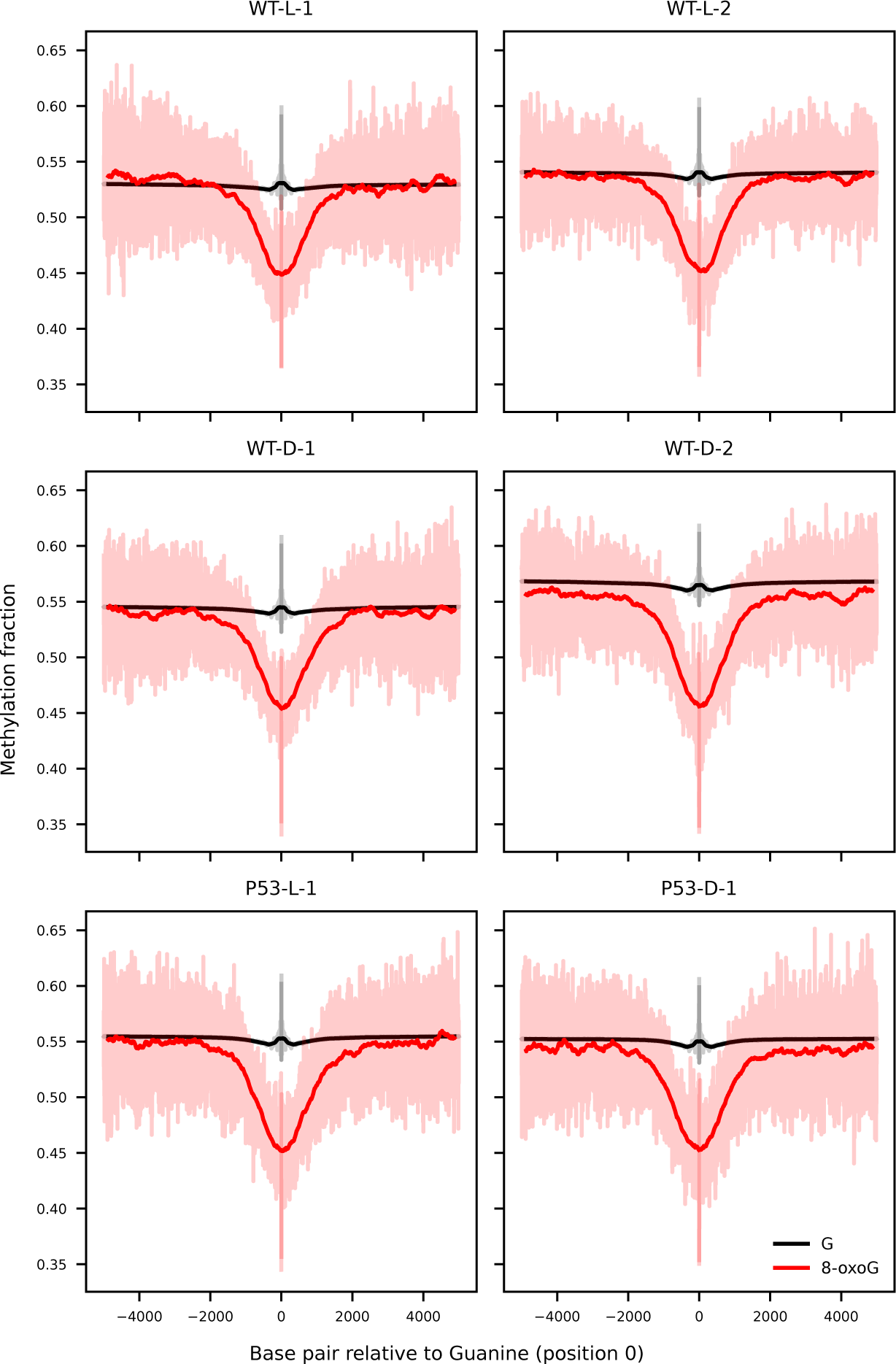
Methylation levels per line relative to 8-oxo-dG. Methylation levels for 8-oxo-dG (red) and Guanine (black) containing reads at position zero, methylation levels are obtained from the same molecule. Transparent colored line indicates the underlying data, the dark line is the result of an 11 base average convolution. Text in the plot indicates the cell line (WT or p53^-/-^), treatment (L-Alanine or D-Alanine), and the biological replicate number.

**Supplementary figure 31:**
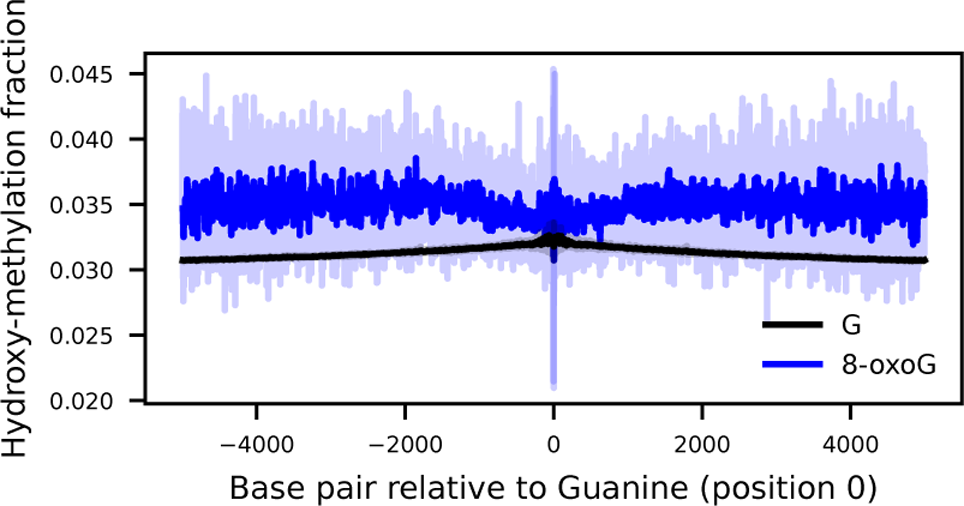
Hydroxy-methylation levels relative to 8-oxo-dG. Hydroxy-methylation levels for 8-oxo-dG (blue) and Guanine (black) containing reads at position zero, hydroxy-methylation levels are obtained from the same molecule. Transparent colored line indicates the underlying data, the dark line is the result of an 11 base average convolution.

**Supplementary figure 32:**
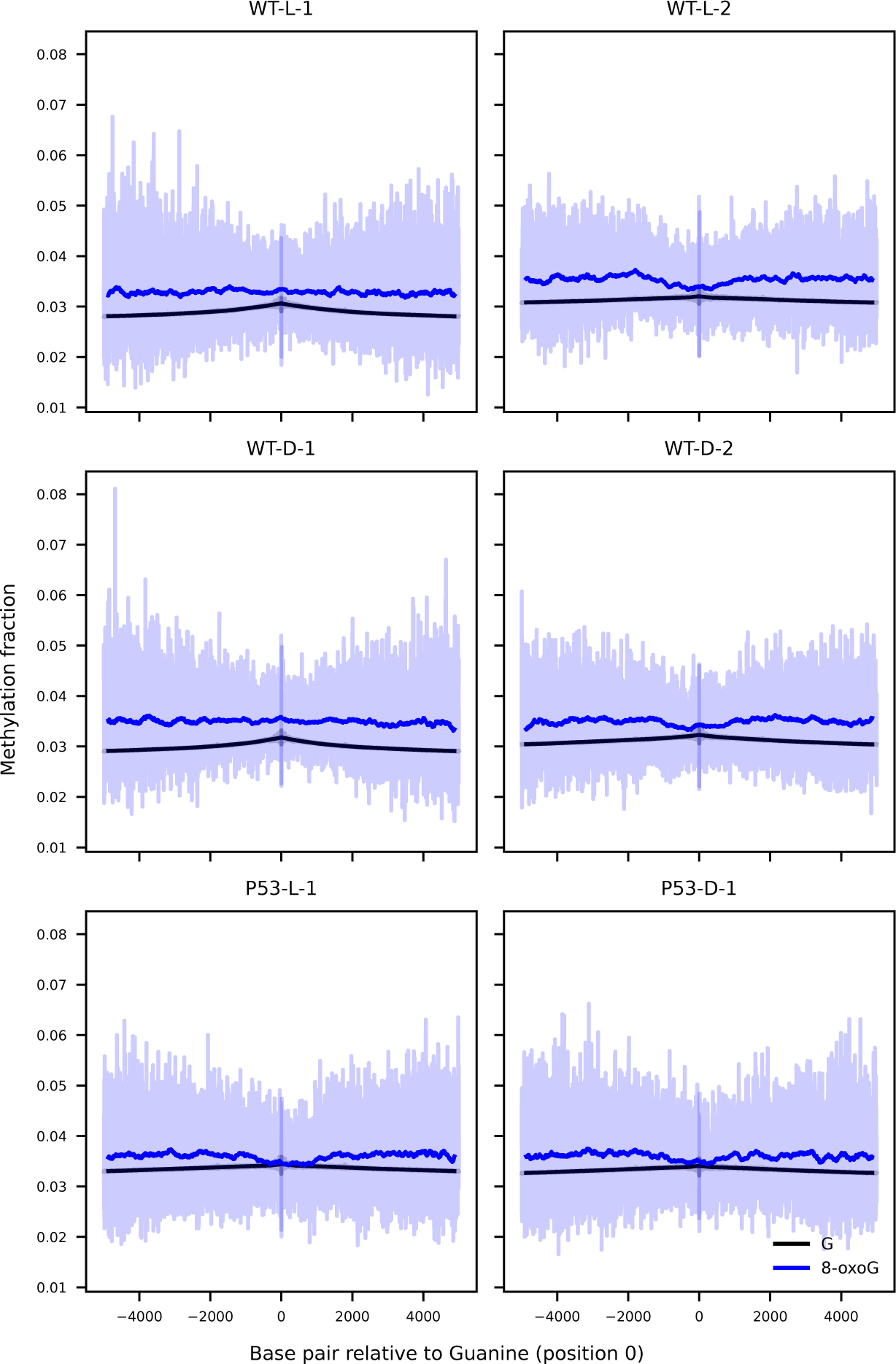
Hydroxy-methylation levels per line relative to 8-oxo-dG. Hydroxy-methylation levels for 8-oxo-dG (blue) and Guanine (black) containing reads at position zero, hydroxy-methylation levels are obtained from the same molecule. Transparent colored line indicates the underlying data, the dark line is the result of an 11 base average convolution. Text in the plot indicates the cell line (WT or p53^-/-^), treatment (L-Alanine or D-Alanine), and the biological replicate number.

**Supplementary figure 33:**
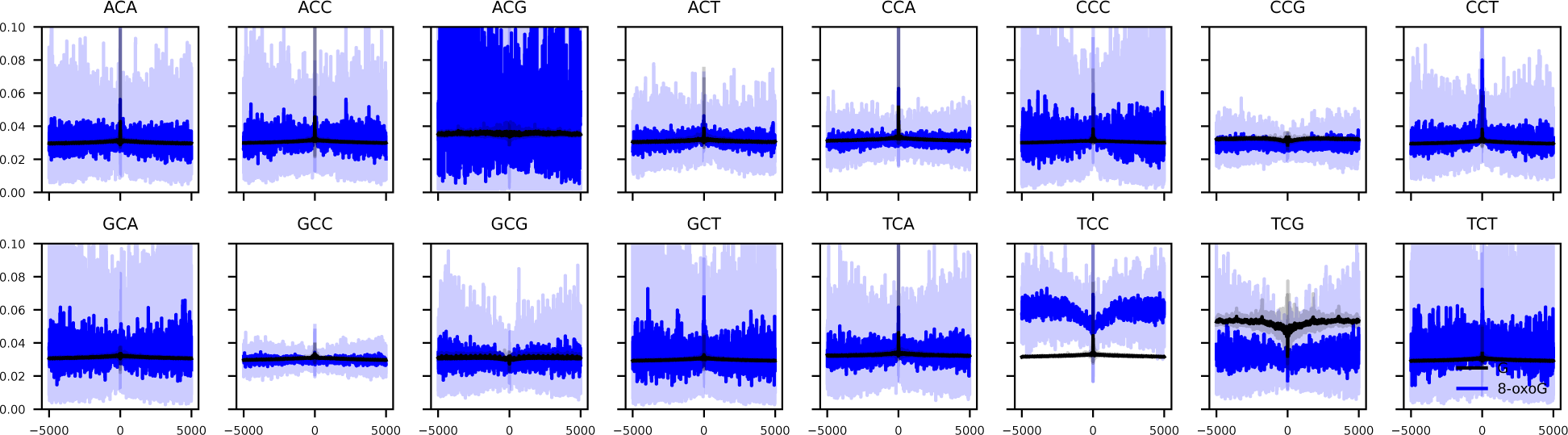
Hydroxy-methylation levels split by 3-mer relative to 8-oxo-dG. Hydroxy-methylation levels for 8-oxo-dG (blue) and Guanine (black) containing reads at position zero, hydroxy-methylation levels are obtained from the same molecule. Data from all conditions is included, and is split between the tri-nucleotide context of the opposite strand of Guanine. Transparent colored line indicates the underlying data, the dark line is the result of an 11 base average convolution.

**Supplementary figure 34:**
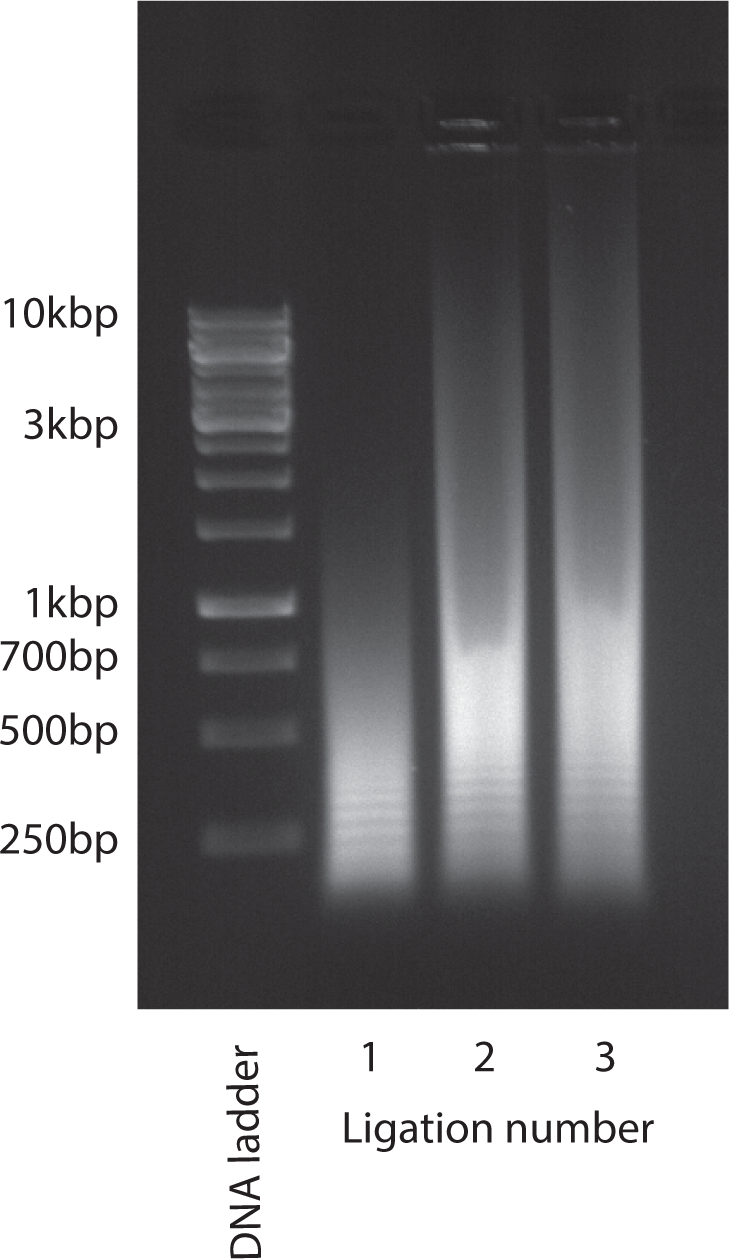
Oligo ligation length. Electrophoresis gel (1% agarose run at 80V for 1 hour) that shows that consecutive repetitive oligo ligations yield increasingly lengthy DNA concatemers.

**Supplementary figure 35:**
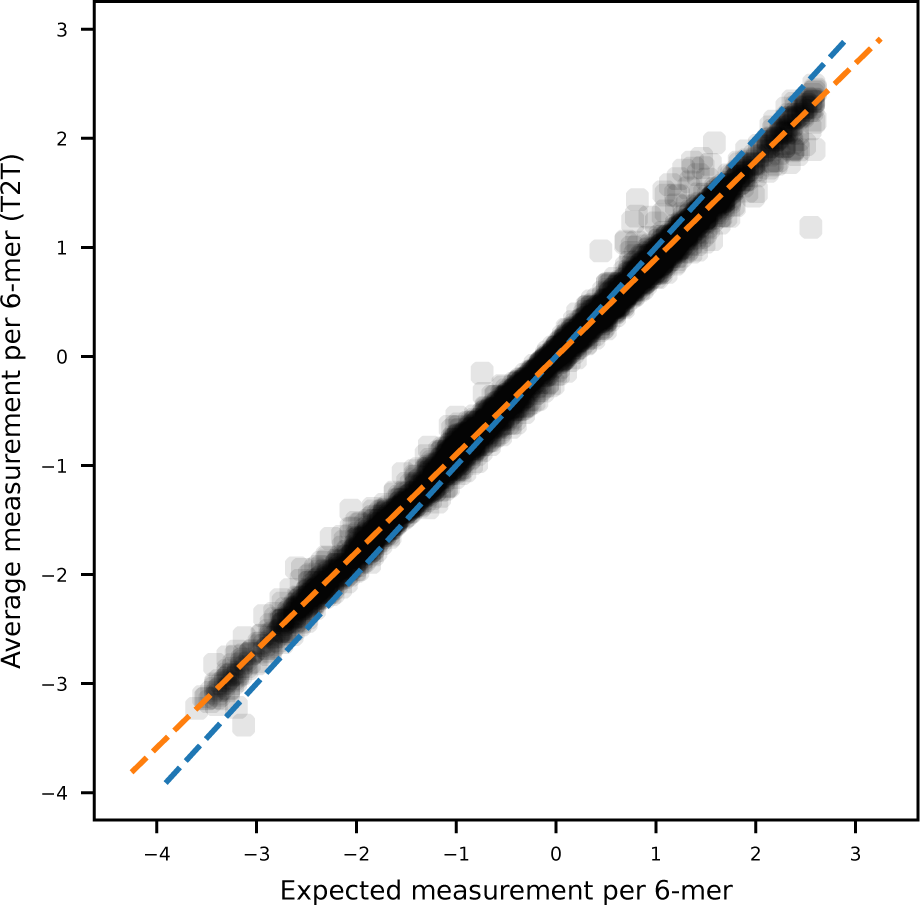
Canonical 6-mer normalization T2T. Measured 6-mer average levels on the T2T dataset before two step signal normalization, compared to expected 6-mer values for non-modified bases. Blue line indicates the identity line. Orange line indicates a linear fit model between the expected and oligo measurements.

**Supplementary figure 36:**
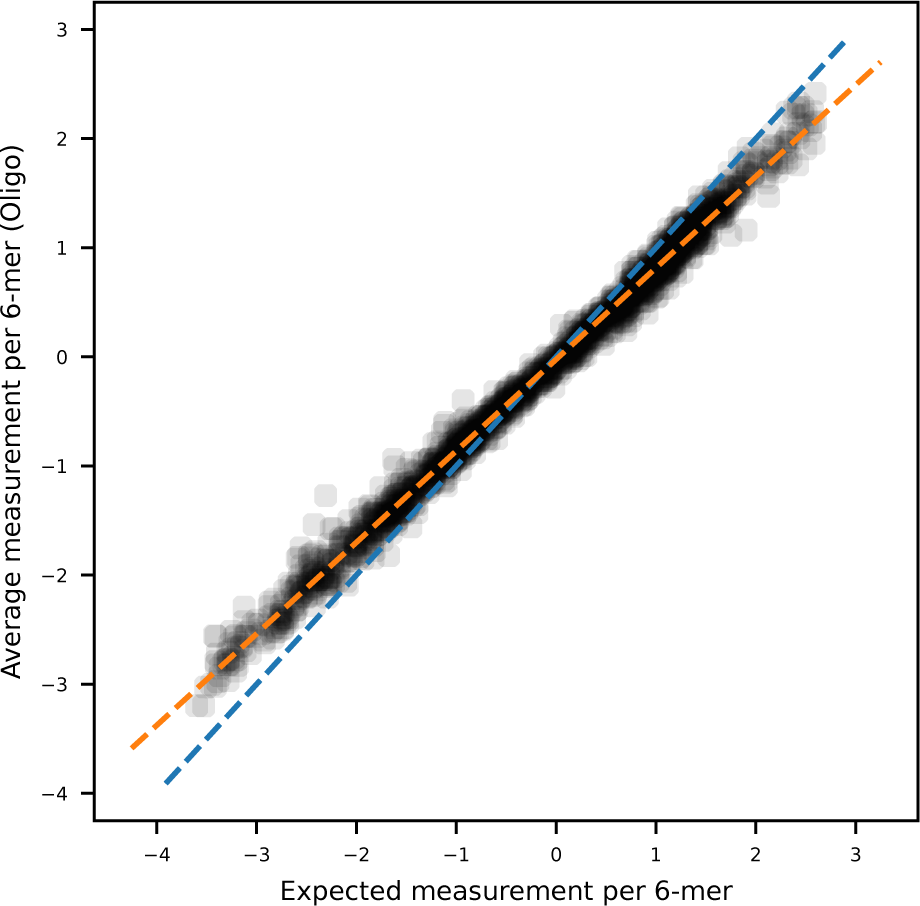
Canonical 6-mer normalization oligo. Measured 6-mer average levels on the oligo dataset before two step signal normalization, compared to expected 6-mer values for non-modified bases. Blue line indicates the identity line. Orange line indicates a linear fit model between the expected and oligo measurements.

**Supplementary figure 37:**
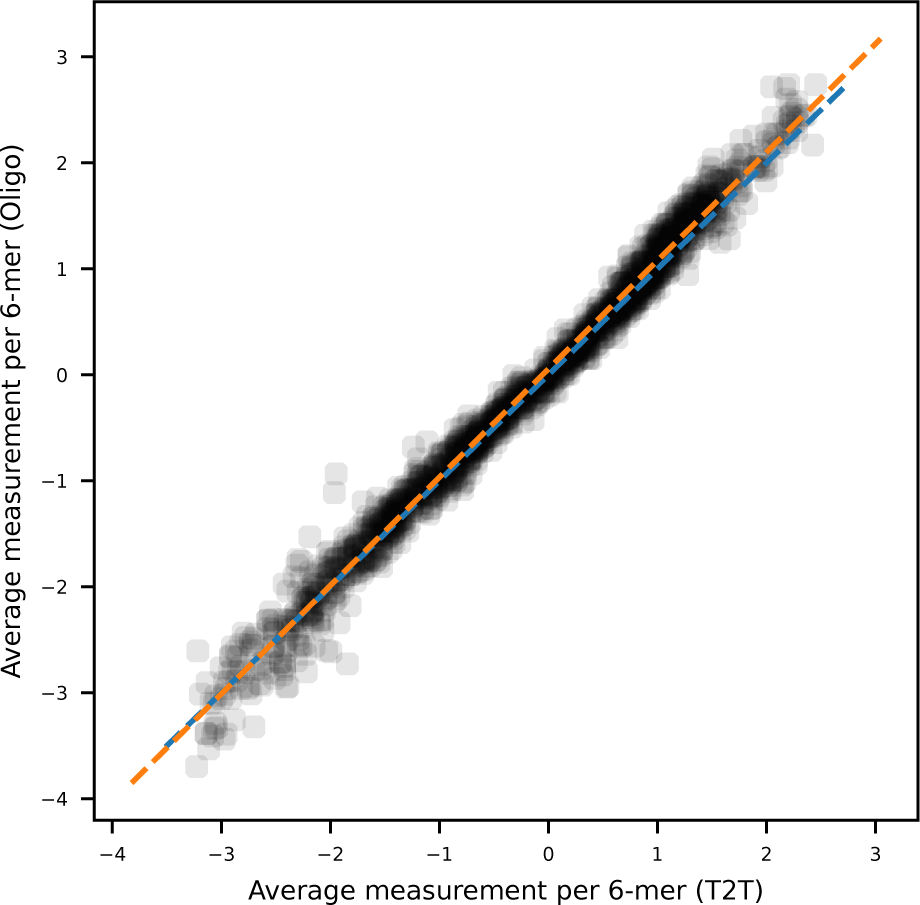
Canonical 6-mer comparison between T2T and oligo. Measured 6-mer average levels on the T2T and oligo datasets after two step signal normalization. Blue line indicates the identity line. Orange line indicates a linear fit model between the T2T and oligo measurements.

**Supplementary figure 38:**
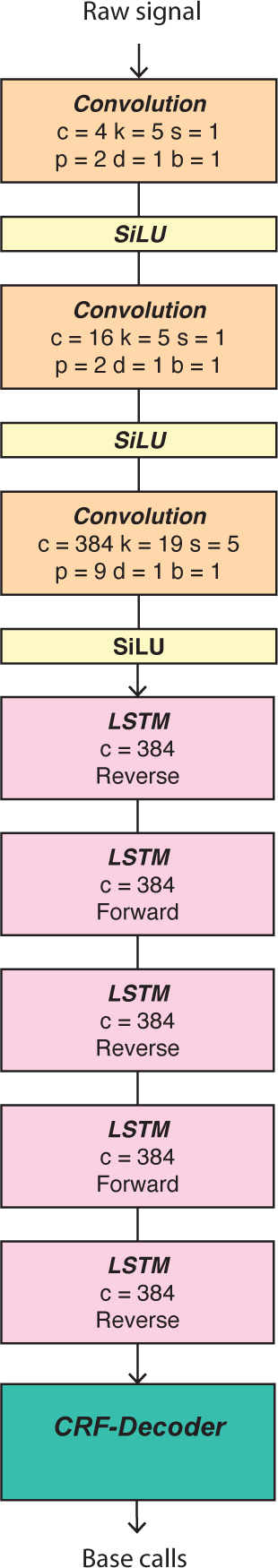
**Neural network architecture of a *Bonito* model.**Schematic representation of the neural network architecture for a *Bonito* model. Numbers indicate output dimension (c), kernel size (k), stride (s), padding (p), dilation (d) and bias (b).

**Supplementary figure 39:**
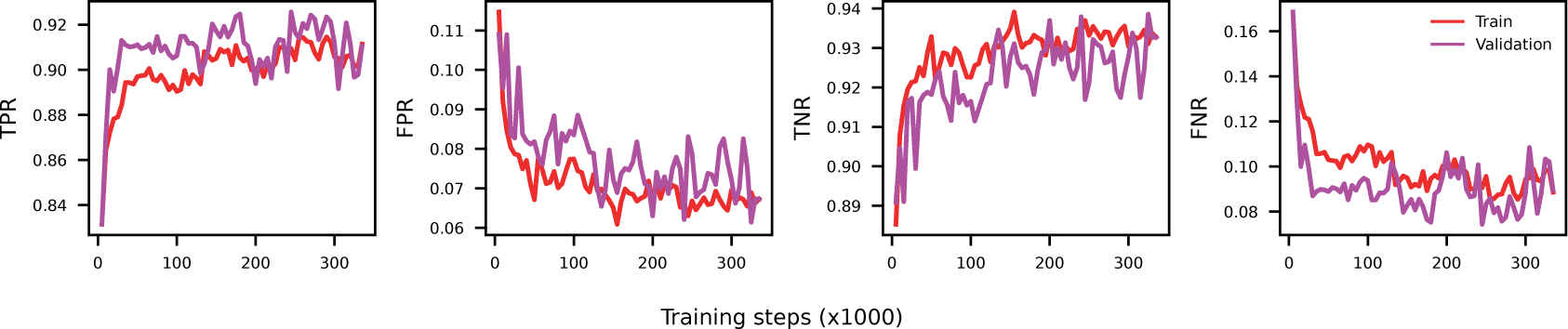
Metrics during training. Performance metrics of the Remora base model during training, trained using a metric learning approach during training. Each plot represents, from left to right, the TPR, FPR, TNR and FNR for the train (red) and validation (purple) folds.

**Supplementary figure 40:**
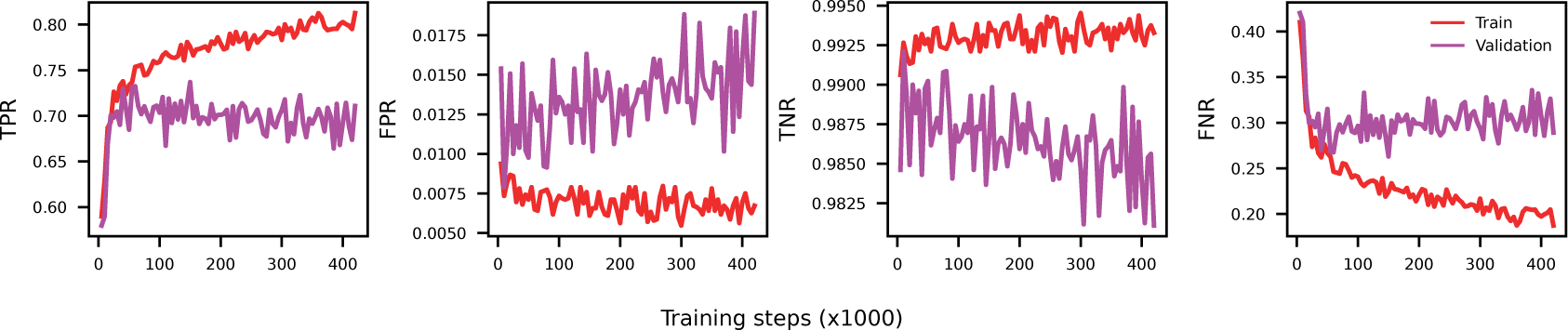
Metrics during training, metric learning. Performance metrics of the Remora base model during training, trained using a metric learning approach during training. Each plot represents, from left to right, the TPR, FPR, TNR and FNR for the train (red) and validation (purple) folds.

**Supplementary figure 41:**
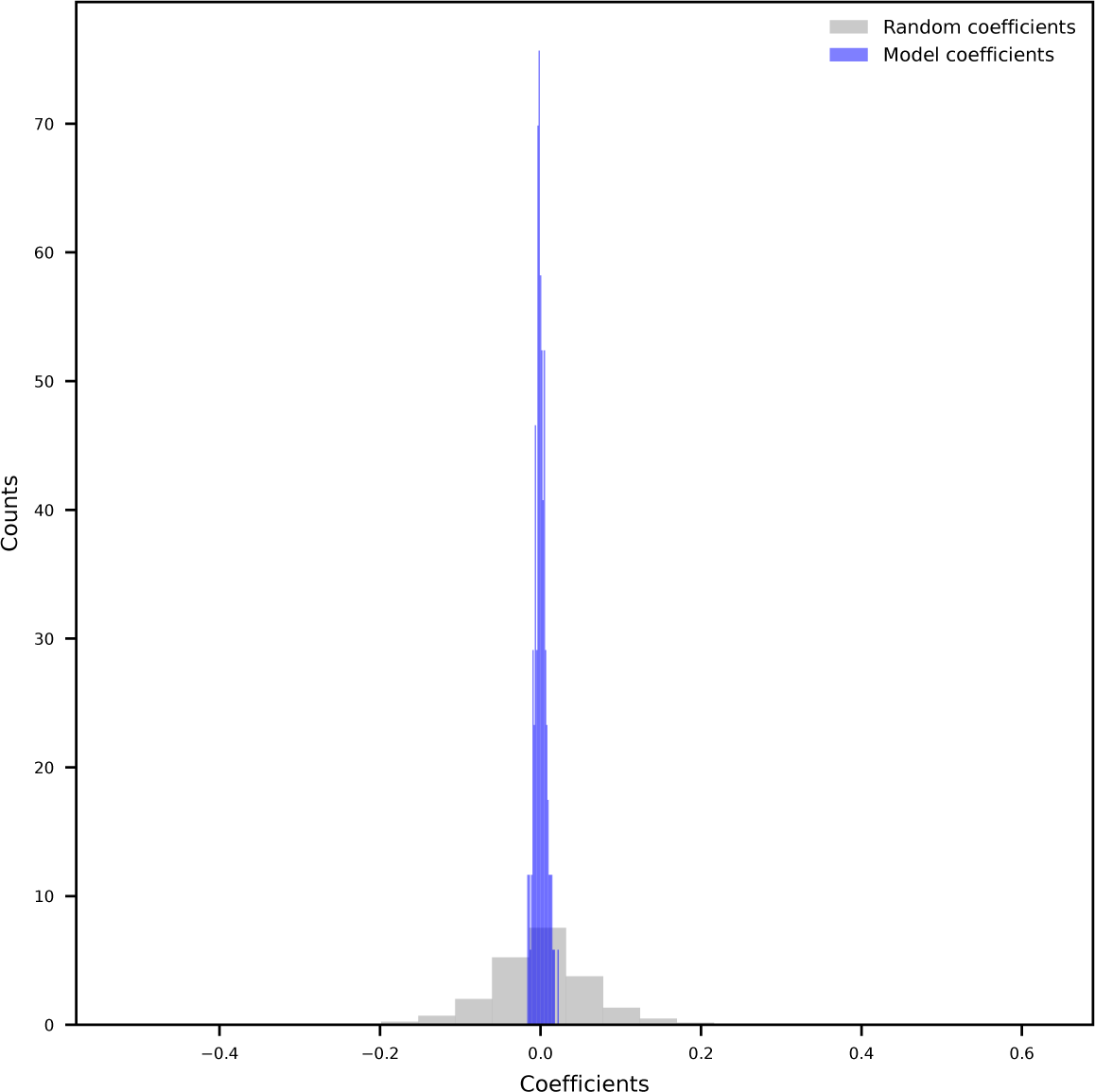
Linear model 5-mer coefficients. Coefficient distribution after fitting a model to explain the observed 8-oxo-dG values per chromosome, chromosome arm and DNA strand based on the 5-mer content. Blue bars indicate the obtained coefficients after model fitting. Gray bars indicate obtained coefficient after label randomization.

**Supplementary figure 42:**
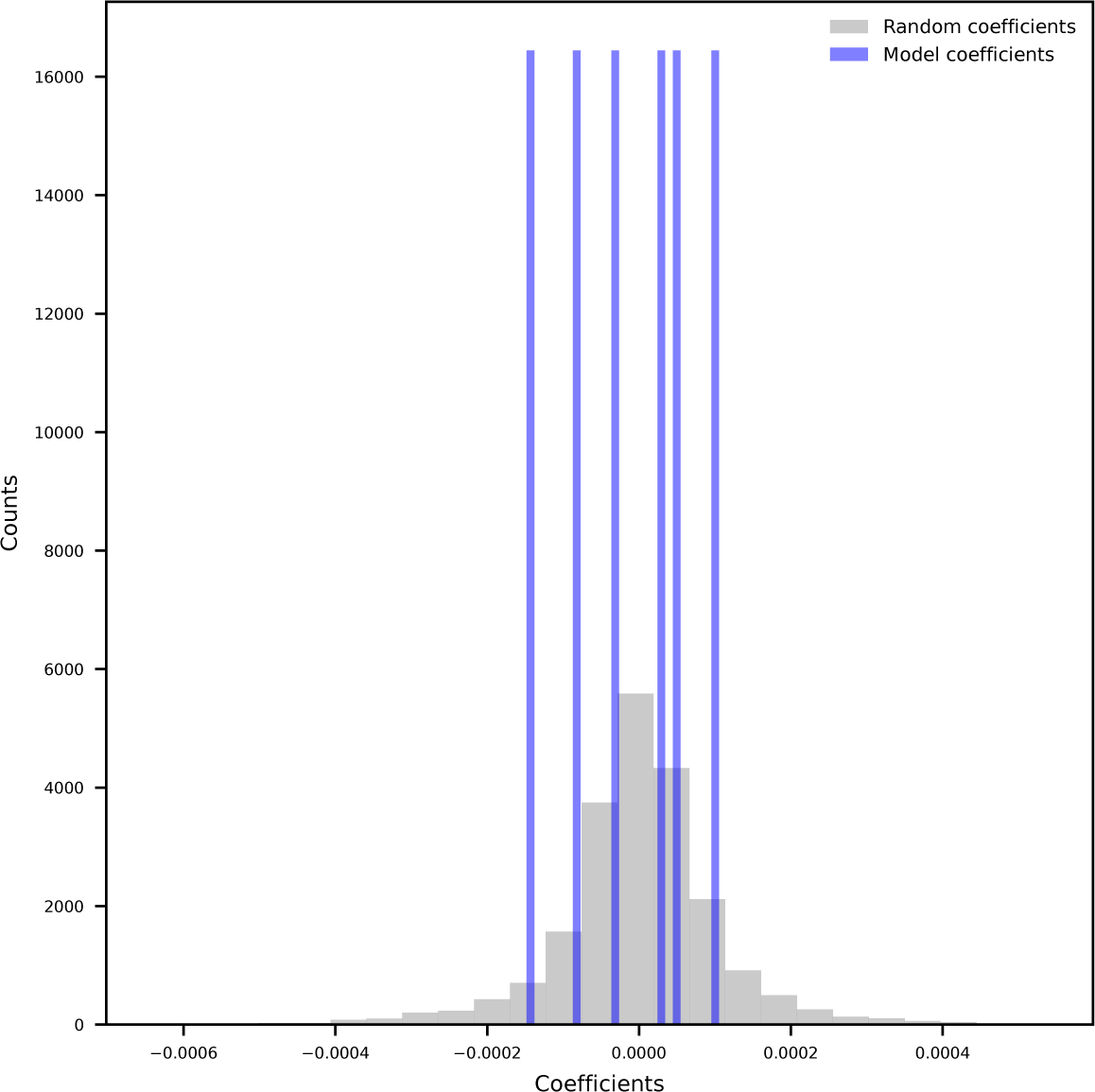
**Linear model genomic regions coefficients.**Coefficient distribution after fitting a model to explain the observed 8-oxo-dG values per chromosome, chromosome arm and DNA strand based on the relative abundance of different genomic regions (exons, introns, intergenic, satellites, centromeres and complex repetitive regions). Blue bars indicate the obtained coefficients after model fitting. Gray bars indicate obtained coefficient after label randomization.

### List of Supplementary Tables

**Supplementary table 1: Oligo sequences.** All synthesized and sequenced oligo sequences. Names indicate the 5-mer in which 8-oxo-dG is. Random bases are indicated as “N” and 8-oxo-dG is indicated as “o”.

**Supplementary table 2: Error rate 8-oxo-dG containing oligos.** Calculated error rate per base in the 8-oxo-dG 5-mer across all sequenced oligos (which contain an 8-oxo-dG). ‘base_3’ corresponds to 8-oxo-dG.

**Supplementary table 3: Error rate non-8-oxo-dG containing oligos.** Calculated error rate per base in the 8-oxo-dG 5-mer across all sequenced oligos (which do not contain an 8-oxo-dG). ‘base_3’ corresponds to cytosine complementary to 8-oxo-dG.

**Supplementary table 4: Performance metrics base *Remora* model.** Performance metrics (TPR, FPR, TNR and FNR) calculated at different thresholds for the trained *Remora* base model, evaluated on the test fold.

**Supplementary table 5: Variant filtering configuration.** Per parameter configuration used for variant filtering for the mutational profile calculation derived from Illumina sequencing.

## Notes

### Summary of Updates

Small text changes, typo corrections, and addition of data upload accession code.

